# Comparative electrostatic mapping of ATP synthase reveals species-dependent axial asymmetry

**DOI:** 10.64898/2026.03.14.711807

**Authors:** Islam K. Matar, Peyman Fahimi, Jean-Nicolas Vigneau, Chérif F. Matta

## Abstract

ATP synthase functions within a highly structured electrostatic environment, but comparative information on its intrinsic protein electrostatics across species remains limited. Here, we analyze 178 crystallographic and cryo-EM ATP synthase structures from 17 species using a consistent Poisson–Boltzmann workflow. The calculations reveal reproducible species-dependent axial electrostatic asymmetries across the F_o_–F_1_ complex, with plane-averaged potential differences of approximately 10–20 mV between an entry-side axial window (*z* = −10 to +10 Å) and an exit-side axial window (*z* = +50 to +70 Å) in several taxa. The five available *Homo sapiens* structures display a consistent axial electrostatic orientation within this species, whereas broader variation is observed across the full multi-species dataset; these two observations reflect within-species and across-species comparisons, respectively. Membrane-mimicking low-dielectric slabs amplify the profiles and demonstrate the sensitivity of the electrostatic landscape to dielectric boundary conditions. We interpret these quantities as static, structure-derived electrostatic descriptors, not as independent membrane voltages or additive proton-motive-force terms. The results provide a comparative electrostatic atlas of ATP synthase and suggest testable hypotheses concerning local proton-pathway energetics, structural evolution, and inhibitor sensitivity.

**GRAPHICAL ABSTRACT:** 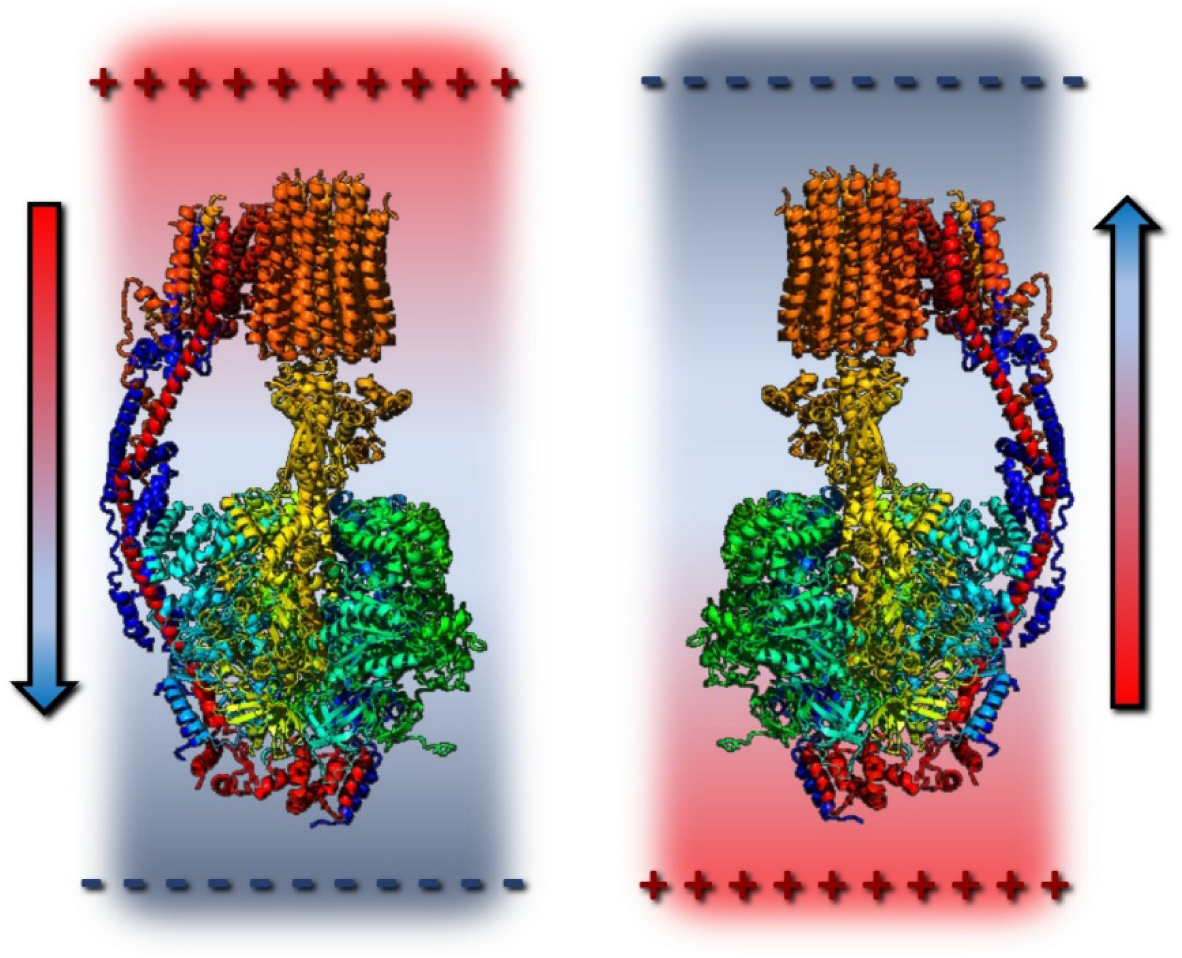

## 1. Introduction

ATP synthase is the rotary enzyme that catalyzes the synthesis of ATP using the proton motive force (pmf) across the mitochondrial, bacterial, or chloroplast’s membrane. The core concept of Mitchell’s chemiosmotic theory [1,2] is that the free energy from oxidative metabolism−stored as an electrochemical proton gradient across the inner mitochondrial membrane, bacterial plasma membrane, and chloroplast thylakoid membrane−is used by the enzyme to drive ATP formation.

During mitochondrial aerobic respiration, electron transport through complexes I– IV of the electron transport chain (ETC) drives protons (H⁺) from the mitochondrial matrix to the intermembrane space (IMS), creating an electrochemical gradient across the inner membrane. This gradient, the proton motive force (pmf) referred to above, has two components: an electrical potential difference (voltage) ΔΨ (matrix negative, IMS positive) and a chemical gradient (difference in proton concentration, often expressed as ΔpH = pH_in_ – pH_out_). The pmf (in units of volts) is defined [3]:

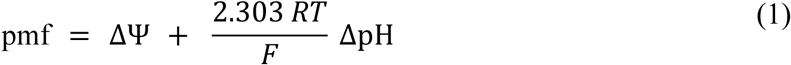

where *R* is the gas constant (8.314 J.mol^―1^.K^―1^), *T* is the absolute temperature, and *F* is Faraday’s constant (96485 C/mol), and 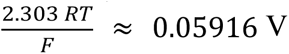 (i.e. ≈ 59 mV/pH unit [3]) at 37 °C for a ΔpH of 1. By convention, ΔΨ is defined as the electrical potential in the interior (e.g. mitochondrial matrix) minus the potential outside (e.g. mitochondrial intermembrane space). This ΔΨ is the electrical component of the transmembrane electrochemical potential (not the full pmf). That is, ΔΨ reflects the net electrical potential difference generated by charge separation across the membrane. In the chemiosmotic expression it is combined with the ΔpH term to define the pmf. Reported mitochondrial ΔΨ ranges from approximately 90 to 225 mV (negative inside) [4] which is actively regulated through an electrical feedback mechanism [5]. For chloroplasts, ΔΨ values as low as −30 mV have been reported, while in bacteria ΔΨ typically ranges from −90 to −180 mV [6]. The relative contributions of ΔΨ and ΔpH to the total pmf vary widely across species and physiological states [6].

The free energy change for a proton moving down its electrochemical gradient (from the IMS back into the mitochondrial matrix) is given by:

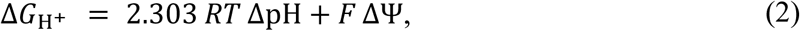

whereby, if we define ΔΨ = Ψ_in_ – Ψ_out_, this formula gives the magnitude of the Gibbs energy gained by a proton moving inward, that is, Δ*G* < 0 indicates a spontaneous flow of protons. Under typical mitochondrial conditions (ΔΨ ≈ −150 to −200 mV and ΔpH ≈ 0.5–1), the proton motive force corresponds to ∼20–30 kJ·mol⁻¹ per proton [3,7]. This energy must be sufficient to overcome the cellular free energy of ATP synthesis.

Although ΔG°′ for ATP synthesis 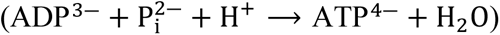 is +30.5 kJ·mol⁻¹ under standard biochemical conditions, the high intracellular [ATP]/[ADP][P_i_] ratio shifts the effective free energy of ATP formation to approximately 50–60 kJ·mol^−1^ under physiological conditions [8–10], which will be referred to as Δ*G*_ATP_. The proton motive force must therefore supply at least this amount of free energy per ATP synthesized, in addition to unavoidable dissipative losses.

In mammalian mitochondria, rotor stoichiometry and metabolite transport together yield an effective requirement of approximately 3.5–4 protons per ATP synthesized and exported to the cytosol [11]. This value reflects both c-ring geometry and the electrogenic cost of ATP/ADP exchange and phosphate import.

ATP synthase is a proton-driven rotary nanomachine in which translocation through the membrane-embedded F_o_ sector generates torque on the c-ring and central γ-shaft, thereby powering ATP synthesis in F_1_ [12]. At a typical mitochondrial proton motive force (≈ 200 mV), each proton delivers ∼0.2 eV [13], yet the intact enzyme rotates at only ∼ 100-150 revolutions per second (rps) under physiological conditions (and up to ∼ 650 rps in thermophilic systems [14]). In contrast, treating the F_o_ c-ring as a quantum mechanical rigid rotor predicts minimum angular velocities (for the first non-zero quantum number (*l* = 1)) on the order of 10^4^–10^5^ rps, one to three orders of magnitude above biological operation [13]. This is consistent with the experiments of Lill *et al.*, who reported a rotation rate of ∼ 4 × 10^4^ rps for isolated F_o_ from chloroplast ATP synthase [15].

Lill et al. report that the “*average*” proton conductance across all isolated F_o_ channels in their experiments is approximately 6,200 H⁺.s⁻¹, a value that includes both active and inactive ATP synthases. However, when considering only the active F_o_ channels (effectively a single-molecule–level analysis based on statistical evaluation), they obtain a conductance of ∼ 6 × 10^5^ H⁺.s⁻¹ per active isolated F_o_ channel, corresponding to ∼ 4 × 10^4^ rps falling exactly within the range of our quantum mechanical calculations [13,15].

Energetically, ATP synthase operates in a strongly overdamped regime in which ∼ 75–83% of proton free energy is captured in ATP and ∼ 17–25% is dissipated, predominantly through internal mechanical friction [13]. Thus, intrinsic electrostatic potentials of the enzyme on the order of 10–20 mV could be relevant if they are sampled locally by protons during translocation through the F_o_ sector. Such potentials are protein-generated electrostatic features that may, in principle, bias local protonation, proton release, proton transfer, or coupling efficiency by approximately 1–2 kJ·mol⁻¹ per proton.

Vigneau *et al*. [16] proposed that ATP synthase carries an intrinsic molecular electrostatic potential (ESP) with a defined orientation along the membrane-normal axis. In the structures examined in that study, the axial ESP was oriented so as to be consistent with a constructive local electrostatic bias for proton translocation [16]. The present work extends that analysis to 178 ATP synthase structures from 17 species and shows that the sign and magnitude of this axial electrostatic asymmetry are species-dependent. In particular, the available human mitochondrial ATP synthase structures exhibit an axial ESP orientation opposite to that observed in several previously examined enzymes. These computed enzyme-linked voltages are clearly static structure-derived electrostatic descriptors that may generate testable hypotheses about local F_o_-sector proton-pathway energetics. We therefore present these computed quantities as static structure-derived axial electrostatic descriptors that may generate testable hypotheses about local F_o_-sector proton-pathway energetics.

## 2. ATP synthase’s intrinsic electrostatic potential (ESP)

Vigneau *et al*. [16] propose that F_o_-F_1_ ATP synthase may possess a protein-intrinsic axial electrostatic potential that could, in principle, influence the local electrostatic environment encountered by translocating protons. Fig. 1 depicts the hypothesis driving the work and that the enzyme’s electric field’s component perpendicular to the membrane plane and its associated electrostatic potential may be parallel or antiparallel to the electrical part of the pmf.

**Fig. 1.**
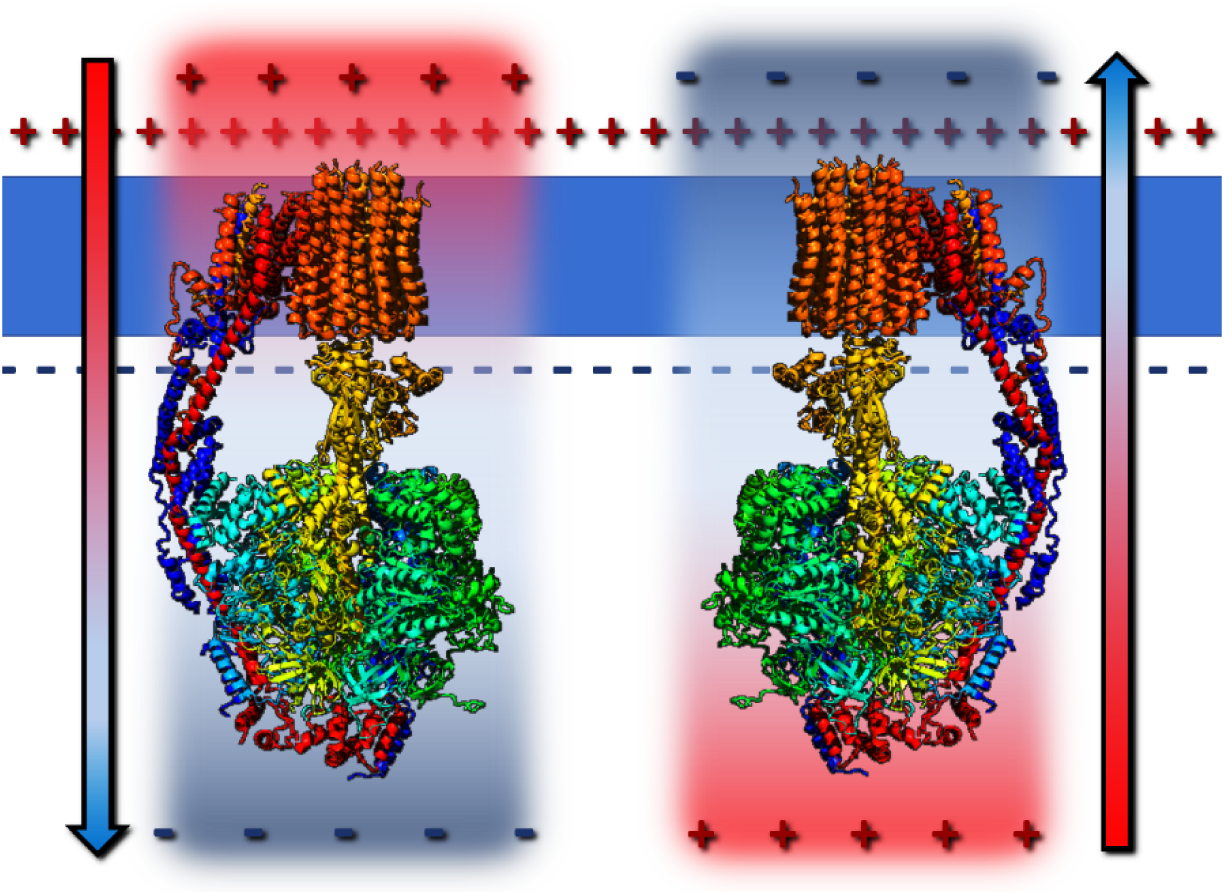
Schematic illustration of the hypothesis: two opposite axial electrostatic polarity classes (axial electrostatic orientation classes) in F_o_-F_1_ ATP synthase: positive-entry axial orientation (*left*) and negative-entry axial orientation (*right*). The membrane bilayer (blue slab) contains the F_o_ sector.

Vigneau *et al*. [16] examined five structures from *Paracoccus denitrificans*, *Bacillus* sp. (bacteria), *S. cerevisiae* (yeast), *Y. lipolytica* (fungus), and *Sus scrofa* (mammal). This analysis is extended to 178 structures retrieved from the Protein Data Bank [17]. Each structure was prepared with PDB2PQR, which was used to assign AMBER force-field atomic partial charges and radii and to generate PQR files [18]. Electrostatic potentials were then computed by numerically solving the finite-difference linearized Poisson–Boltzmann equation (LPB) using the Adaptive Poisson–Boltzmann Solver (APBS) [19] with pH = 7.0, T = 298 K (25 °C), ε(solvent) = 78.5, ε(protein interior) = 6, and where applicable, ε(membrane) = 4. In all calculations, a bulk ionic strength of 150 mM NaCl was maintained [20]. Calculated electrostatic potential grids surrounding each structure were analyzed using an in-house Nextflow/Python pipeline (see Methodology). The linearized formulation of the PB equation is formally valid in the regime ∣*eΨ*∣ ≪ *k*_B_*T* (i.e., when ∣Ψ∣≪∼25 mV at physiological temperature) since the nonlinear ion-density term sinh (*e*Ψ/*k*_B_*T*) may be approximated to first order [20–22]. Under physiological screening conditions (Debye length ≈ 8 Å at 150 mM – *vide infra*), nonlinear corrections are reduced by ionic screening, and LPB provides an accurate first-order description of biomolecular electrostatics for comparative analyses. Importantly, the linearized equation yields a linear elliptic partial differential equation that can be solved *via* a single sparse linear system, avoiding the iterative nonlinear solution procedures required by the full Poisson–Boltzmann equation. This substantially improves numerical robustness and reduces computational cost for large-scale, cross-species comparison (178 structures) under identical solvent conditions. Because all structures were treated consistently within the same dielectric and ionic framework, the LPB formulation provides a stable and internally consistent basis for comparative electrostatic analysis. Because all structures were treated under identical dielectric and ionic conditions (150 mM NaCl), any systematic nonlinear effects would apply uniformly across the dataset and are therefore unlikely to alter the comparative trends or relative ordering reported here.

All structures studied in the past [16] exhibit constructive contributions from the ESP of ATP synthase (ΔΨ_ATP synthase_) to the pmf. As an example, see the Ψ_ATP synthase_ of fungus *Yarrowia lipolytica*, PDB Code # 5FL7, in Fig. 2. ATP synthase’s plane-averaged electrostatic potential (⟨Ψ_ATP synthase_(*x*,*y*|*z*)⟩) of this structure exhibits a net drop along the *z*-axis (pointing from the intermembrane-space perpendicularly toward the mitochondrial matrix). The average ESP of this fungus is positive near the upper vestibule and becomes progressively more negative along the proton pathway toward the matrix. The (average) axial voltage between the point of entry and exit of the proton into and from the membrane domain, i.e., the enzyme-dependent voltage, is defined:

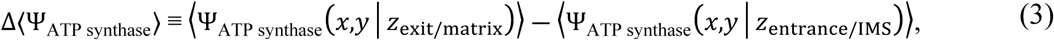

where ΔΨ_ATP synthase_ is defined analogously to the chemiosmotic membrane potential as Ψ_matrix_ − Ψ_IMS_ (given the convention that Ψ_matrix_ < Ψ_IMS_ under normal oxidative phosphorylation). For clarity, Δ〈Ψ_ATP synthase_〉 means the difference of the averages and not the average of the differences.

**Fig. 2.**
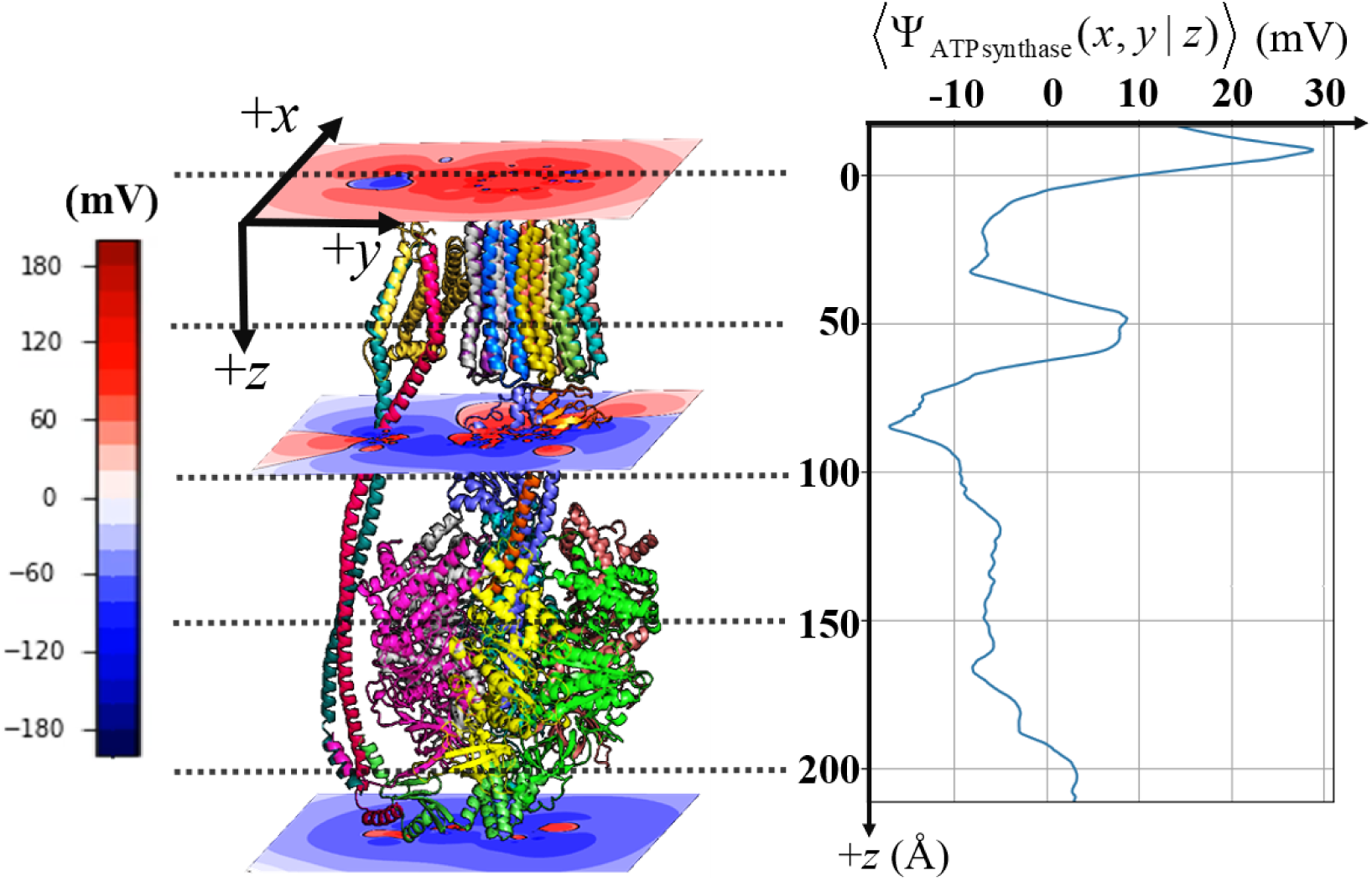
Average electrostatic potential of ATP synthase of *Yarrowia lipolytica* (PDB Code # 5FL7) per cross-sectional *xy*-planes as a function of the *z*-coordinate (the long axis of the protein perpendicular to the inner mitochondrial membrane). ***Left:*** The enzyme is shown with three representative *xy*-slices colored by electrostatic potential (red, positive; blue, negative) -with the color-scale showing the values of the electrostatic potential in mV. A right-handed Cartesian coordinates is indicated with +*z* increasing downward. Dotted lines mark the approximate *z*-positions of the slices. ***Right:*** Profile of the average potential per *xy*-plane, ⟨ΔΨ_ATP synthase_(*x*,*y*|*z*)⟩ in mV plotted against *z* in Å using the same +*z*-down convention (horizontal dotted lines roughly correspond to the slice planes on the left). This curve summarizes how the mean electrostatic environment varies along the axis perpendicular to the membrane.

The proton translocation path “reaction” is written as:

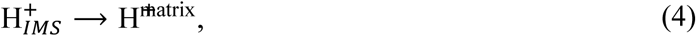

where IMS = intermembrane space, and where the angular brackets in Eq. (3) indicate averaging over an *xy*-plane (see Methods), and where *z*_IMS_ and *z*_matrix_ are the approximate *z*-coordinates on the intermembrane space side and on the matrix side, respectively. In bacteria, “entrance/intermembrane space” and “exit/matrix” in Eqs. (3) and (4) correspond to the extracellular medium and the cytosol, respectively. In the case of the five previously studied organisms [16], ΔΨ_ATP synthase_ has the same sign as the transmembrane potential (ΔΨ_membrane_) reinforcing the electric component of the pmf.

The enzyme-linked electrostatic potentials reported by Vigneau *et al*. correspond to energy shifts of approximately 1–3 kJ·mol^−1^ per proton. Given that cellular ATP synthesis requires ∼ 50–60 kJ·mol^−1^ per ATP and ∼ 3–4 protons per ATP in mammalian mitochondria (depending on c-ring stoichiometry and transport costs), the cumulative effect over oxidative phosphorylation can be non-negligible. Because Δ*G*_ATP_, proton stoichiometry, and ATP yield per substrate molecule vary with organism, metabolic state, and carbon source [23,24], the resulting correction to the total bioenergetic budget is system-dependent. Under representative mitochondrial conditions, however, a 1–3 kJ·mol^−1^ shift per proton can plausibly alter the effective energetic balance by several percent to at most of the order of ∼ 10%.

The publication of the structures of the human enzyme [25,26] has prompted the extension of this work to also enlarge the comparison set. Hence, 178 electron microscopy and electron diffraction structures of ATP synthase from 17 different organisms (including *Homo sapiens*) were obtained from the Protein Data Bank [17], including the five previously studied structures. This dataset includes structures of 17 species which, by kingdom, include ***Animalia*** (mitochondrial) [*Bos taurus* (12 structures), *Homo sapiens* (5), *Ovis aries* (1), and *Sus scrofa* (2)]; ***Bacteria*** (plasma membrane bound) [*Acinetobacter baumannii* (3 structures), *Bacillus sp.* (3), *Escherichia coli* (20), *Mycobacterium tuberculosis* (3), *Mycolicibacterium smegmatis* (24), and *Paracoccus denitrificans* (1)]; ***Fungi*** (mitochondria) [*Ogataea angusta* (3 structures), *Saccharomyces cerevisiae* (39), and *Yarrowia lipolytica* (1)]; ***Plantae*** (chloroplast membrane bound) [*Polytomella sp.* (45 structures) and *Spinacia oleracea* (12)]; and ***Protista*** [*Euglena gracilis* (mitochondria) (3 structures) and *Toxoplasma gondii* (mitochondrial) (1)]. These groups do not yet establish a direct phylogenetic or physiological rule. They suggest, however, that ATP synthase electrostatics may vary with membrane type, c-ring composition, lipid environment, respiratory strategy, and pmf partitioning. The species-dependent classes identified here are to be taken as *comparative electrostatic phenotypes* rather than as functional categories. Whether similar electrostatic descriptors cluster along specific evolutionary branches, membrane types, or metabolic lifestyles will require a larger phylogenetically controlled dataset and direct experimental measurements of pmf, proton flux, and ATP synthesis; this is beyond the scope of the present study.

All 178 structures were subjected to the same protocol mentioned above (see “Methods”) whereby *the entire protein is solvated in an aqueous solution of NaCl of 150 mM without any explicit membrane region*. Hence, ΔΨ_ATP synthase_ values obtained in full aqueous solvation represent lower bounds for values if the membrane’s much lower dielectric constant was accounted for (which would magnify the electric potential and field).

The results of the enzyme-dependent voltage (ΔΨ_ATP synthase_) are shown in Figs. 3 and 4. Fig. 3 overlays the 178 plane-averaged electrostatic potential profiles 〈Ψ_ATP synthase_〉 along the enzyme’s long *z*-axis (Figs. 1–2). Structures were classified according to the plane-averaged electrostatic-potential profile in the membrane-proximal entry-side window, *z* = −20 to +20 Å. The classification was based on the first prominent peak detected in this window. When no clear peak was detected, the mean plane-averaged potential over the same window was used instead. Technical and coding details are provided in the Supporting Information. This window was chosen to represent the electrostatic environment encountered by a proton as it approaches the F_o_ sector during ATP synthesis. On this basis, three groups emerge: positive-entry profiles (⟨Ψ⟩ > +10 mV; 131 structures), ambiguous-moderate profiles (0 to +10 mV; 12 structures), and negative-entry profiles (⟨Ψ⟩ < 0 mV; 35 structures, including all five human structures). The classification into three categories in Fig. 3 was therefore based on this finite averaging window. A small number of borderline cases lie near the threshold between categories; minor smoothing or structural differences may therefore make some curves appear visually closer to an adjacent panel without altering the sign-based grouping criterion. This classification is therefore descriptor-based. The groups are defined by the sign and magnitude of the plane-averaged entry-side electrostatic potential in a fixed axial window, applied uniformly to all structures. This differs from recent machine-learning and multidimensional embedding approaches to protein electrostatic classification, which compare higher-dimensional electrostatic features across protein sets [32]. Our aim here is narrower being limited to define an ATP-synthase-specific axial electrostatic descriptor along the membrane-normal direction. Because no independent experimental labels are currently available for constructive versus opposing ATP-synthase electrostatic function, supervised classification was not used in the present study.

**Fig. 3.**
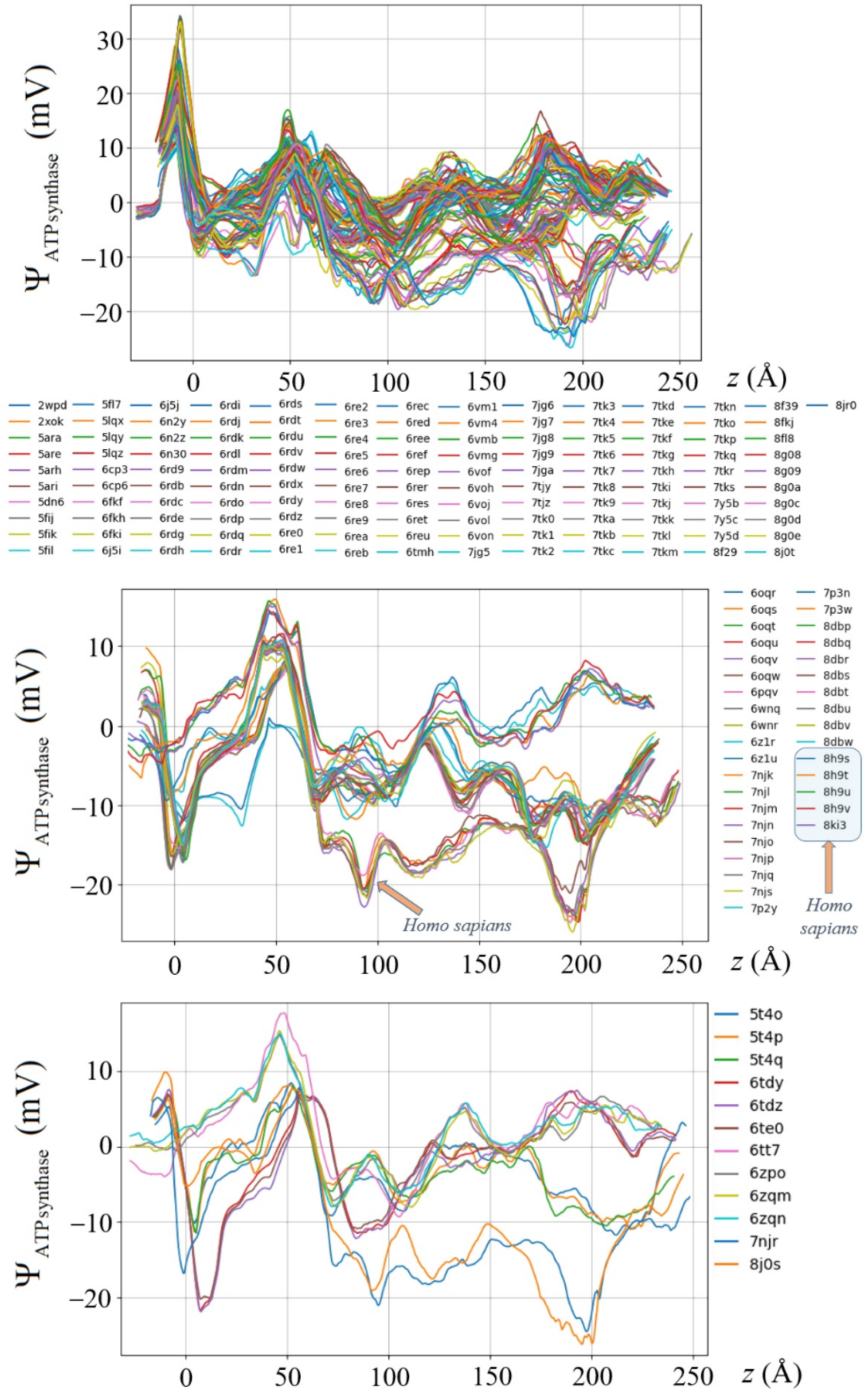
Axial plane-averaged electrostatic potentials for the 178 ATP synthase structures (PDB codes in legend). Each trace shows ⟨Ψ(*x*,*y*|*z*)⟩ (mV), obtained by averaging the electrostatic potential over planes perpendicular to the molecular long (*z*-) axis. Profiles are grouped into three panels according to their low-*z* (IMS-side) behavior: **(top)** initial positive band; **(middle)** ambiguous: near-zero to moderately negative or positive entry; **(bottom)** pronounced negative trough at entry. All calculations were performed in bulk aqueous medium (ε = 78.5) with 150 mM NaCl and no explicit membrane; thus, the absolute magnitudes represent lower bounds relative to membrane-embedded models. The five *Homo sapiens* structures (indicated in the legend) cluster within the negative-entry group and exhibit the same sign and orientation of the axial potential gradient at the IMS interface (i.e., a reproducible negative potential upon entry from low *z*, followed by recovery toward positive values). For clarity, only 10 distinct line colors are used in the main figure; individual traces for all 178 structures are provided in the Supplementary Information.

**Fig. 4.**
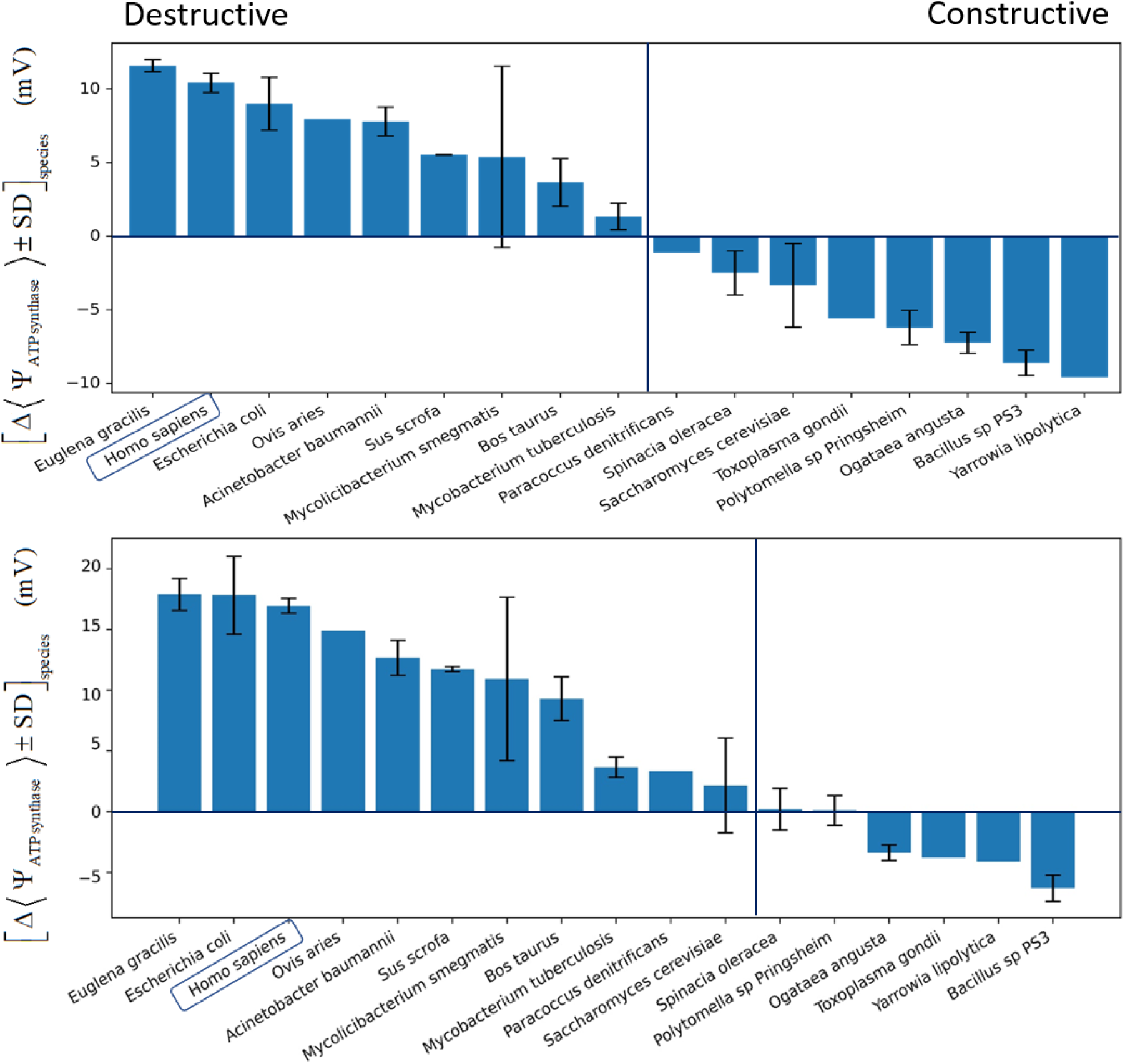
Species-dependent enzyme-linked voltage in aqueous and membrane-embedded models. **Top:** ATP synthase–dependent potential difference (ΔΨ_ATP synthase_) computed in full aqueous medium (150 mM NaCl) for 17 species calculated as the difference between the mean potential difference at proton entry (*z* = −10 to +10 Å) and exit (z = +50 to +70 Å) windows along the enzyme’s long axis. **Bottom:** Corresponding ΔΨ_ATP synthase_ values after embedding the F_o_ sector in a planar dielectric slab (ε = 4) to mimic the membrane core. The slab extends from *z* = +10 to +50 Å. In both bar-graphs, “destructive” (negative-entry) and “constructive” (positive-entry) are defined according to their opposition or alignment to the physiological ΔΨ_ATP synthase_ sign, respectively (Eq. (3)). Comparison of the two bar graphs isolates the influence of membrane dielectric effects on the enzyme-linked voltage (see also Fig. 5).

The axial electrostatic asymmetry descriptor ΔΨ_ATP synthase_ was calculated by averaging the ESP over a 20 Å window on each side of the membrane to compare species based on the potential difference. The ESP was averaged from *z* = −10 to +10 Å on the entry-side axial window and from *z* = +50 to +70 Å on the exit-side axial window, and ΔΨ_ATP synthase_ is defined as the difference between these two averages. The same procedure was applied to each structure and the resulting values were averaged across all structures available for each species. The resulting species means ± SD are presented in Fig. 4.

To quantify the potential difference experienced by a proton during translocation, we define an enzyme-linked voltage, ΔΨ_ATP synthase_, as the difference between the mean plane-averaged electrostatic potentials in two axial windows flanking the membrane. Specifically, the potential was averaged over *z* = −10 to +10 Å (entry-side axial window) and *z* = +50 to +70 Å (exit-side axial window), and their difference was taken as ΔΨ_ATP synthase_. For each species, values were averaged over all available structures (Fig. 4).

In fully aqueous medium (ε = 78.5, 150 mM NaCl; Fig. 4a), ΔΨ_ATP synthase_ is typically modest (within ±10 mV), indicating that the intrinsic protein electrostatics alone generate a small axial voltage bias. In our previous preliminary work [16] we measured the potential difference between two points at the beginning and end of the c-ring, while in this work we average over 20 Å at both sides of the membrane which is why our numerical values here are more reliable (but the trends for the five structures remains intact). In contrast, embedding the F_o_ sector in a membrane-mimicking dielectric slab (ε = 4, spanning *z* = +10 to +50 Å; Fig. 4b) slightly amplifies the magnitude of ΔΨ_ATP synthase_ (spanning 21.2 mV without the membrane to 24.2 mV with the membrane representing slab) and alters its species-dependent polarity (axial electrostatic orientation). This comparison isolates the dielectric focusing effect of the membrane, i.e., compression of electric field lines within the low-dielectric slab at the membrane–water interfaces.

Although the insertion of a low-dielectric slab (ε = 4) generally amplifies the axial field through dielectric focusing, it *does not act as a uniform multiplicative rescaling of the enzyme-linked voltage* when 150 mM NaCl is included. Even under the linearized Poisson–Boltzmann approximation adopted in this work, which in SI units is expressed as:

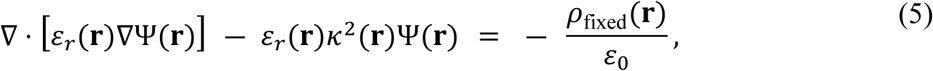

where

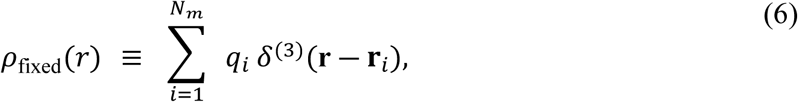

*ε_r_*(**r**) is the relative dielectric constant, *δ*^(3)^ is a three-dimensional Dirac delta function (with units of m⁻^3^), and where the screening term *κ*²Ψ operates only within ion-accessible solvent regions (*κ* = 0 inside the membrane slab since there are no mobile ions) and depends on the Debye length associated with the salt concentration. The slab representing the membrane alters the spatial dielectric map *ε_r_*(**r**) and the geometry of ion-accessible space, thereby modifying how the screening operator and dielectric boundary conditions interact with each enzyme’s specific fixed charge distribution. Because the solution remains sensitive to the detailed spatial arrangement of charges and dielectric discontinuities, the change from fully aqueous to slab-embedded conditions can perturb the window-averaged axial potentials in a structure-dependent manner. Consequently, rank ordering of ΔΨ_ATP synthase_ across species need not be preserved, even though the underlying equation is linear. In contrast, in vacuum or in salt-free water, one would expect the slab primarily to produce a more uniform dielectric focusing effect with greater preservation of ordering [20,21,27–29].

The increase in ΔΨ_ATP synthase_ upon membrane embedding is substantially smaller than it would be in the absence of electrolyte. In salt-free water (κ = 0), the low-dielectric slab would primarily act through dielectric focusing, increasing the axial potential difference without competing screening effects. At 150 mM NaCl, however, the linearized Poisson– Boltzmann term introduces Debye screening with a characteristic decay length of ∼ 8 Å. (At 298 K, the Debye screening length for a 1:1 electrolyte is *λ*_D_ = [(ε₀ε*k*_B_*T*/(2*N*_A_*e*²*I*)]^½^; substituting *ε* ≈ 78.5 and *I* = 0.150 M gives *λ*_D_ ≈ 0.78 nm [30,31]), so electrostatic fields are exponentially attenuated over nanometer scales in the solvent.). As a result, long-range electrostatic fields are exponentially attenuated in the solvent, and part of the dielectric focusing enhancement is offset by ionic screening. The net effect is a moderated increase in ΔΨ_ATP synthase_ relative to a vacuum or salt-free aqueous calculation. This attenuation is an expected consequence of electrolyte screening rather than a limitation of the slab model.

Notably, the five *Homo sapiens* structures available in the dataset cluster within the negative-entry group, indicating within-species consistency among the currently available human structures. The human enzyme remains consistently in the destructive regime upon membrane embedding.

Because ΔΨ_ATP synthase_ is approximately zero in *M. tuberculosis* but negative (with larger uncertainty) in *M. smegmatis* in aqueous conditions, the latter may experience a slightly reduced effective electrical driving component under identical membrane potentials. While speculative, such electrostatic differences could influence sensitivity to oxidative phosphorylation inhibitors that collapse the proton-motive force (e.g., diarylquinolines), and therefore warrant experimental examination. The mycobacterial results should be interpreted cautiously but may be relevant to ATP-synthase-directed drug discovery. *M. tuberculosis* shows a near-zero axial electrostatic asymmetry, whereas *M. smegmatis* trends more negative and is more variable. Because *M. smegmatis* is frequently used as a surrogate for mycobacterial screening, this difference suggests that responses to oxidative-phosphorylation inhibitors, including diarylquinolines that target the F_o_/c-ring region [33], may not transfer quantitatively between the two species. The present results do not establish altered drug sensitivity, but they identify a structural-electrostatic variable that should be considered when interpreting surrogate-model screens and should be tested by early confirmation in *M. tuberculosis* using ATP, membrane-potential, oxygen-consumption, and survival assays.

Fig. 5 provides the underlying one-dimensional axial electrostatic profiles from which the human bars in Fig. 4 are derived, using two representative structures: 8H9T (without inhibitor) and 8H9S (with endogenous 106-residue F_1_F(_o_)-ATPase inhibitor [25] which prevents ATP hydrolysis when membrane potential collapses and the enzyme reverses, thereby preserving cellular ATP under depolarizing conditions [34]). In bulk solvent (left panels), the axial potential difference between the two membrane-side windows fluctuates modestly (≈ ±10–15 mV), consistent with Fig. 4a. After insertion into the low-dielectric slab (right panels), the range of potential increases sharply within the slab region, exceeding 200 mV before decaying outside the membrane boundaries. The difference between the same axial windows now yields the amplified values shown for *Homo sapiens* in Fig. 4b. Comparison of 8H9T and 8H9S shows that the inhibitor produces only minor perturbations to the axial electrostatic profile relative to the dominant dielectric amplification introduced by the membrane. This explains the small standard deviation observed for the human group in Fig. 4. This comparison provides an initial test of state sensitivity for the human enzyme. A systematic ATP/ADP-state comparison across matched structures remains an important future extension.

**Fig. 5.**
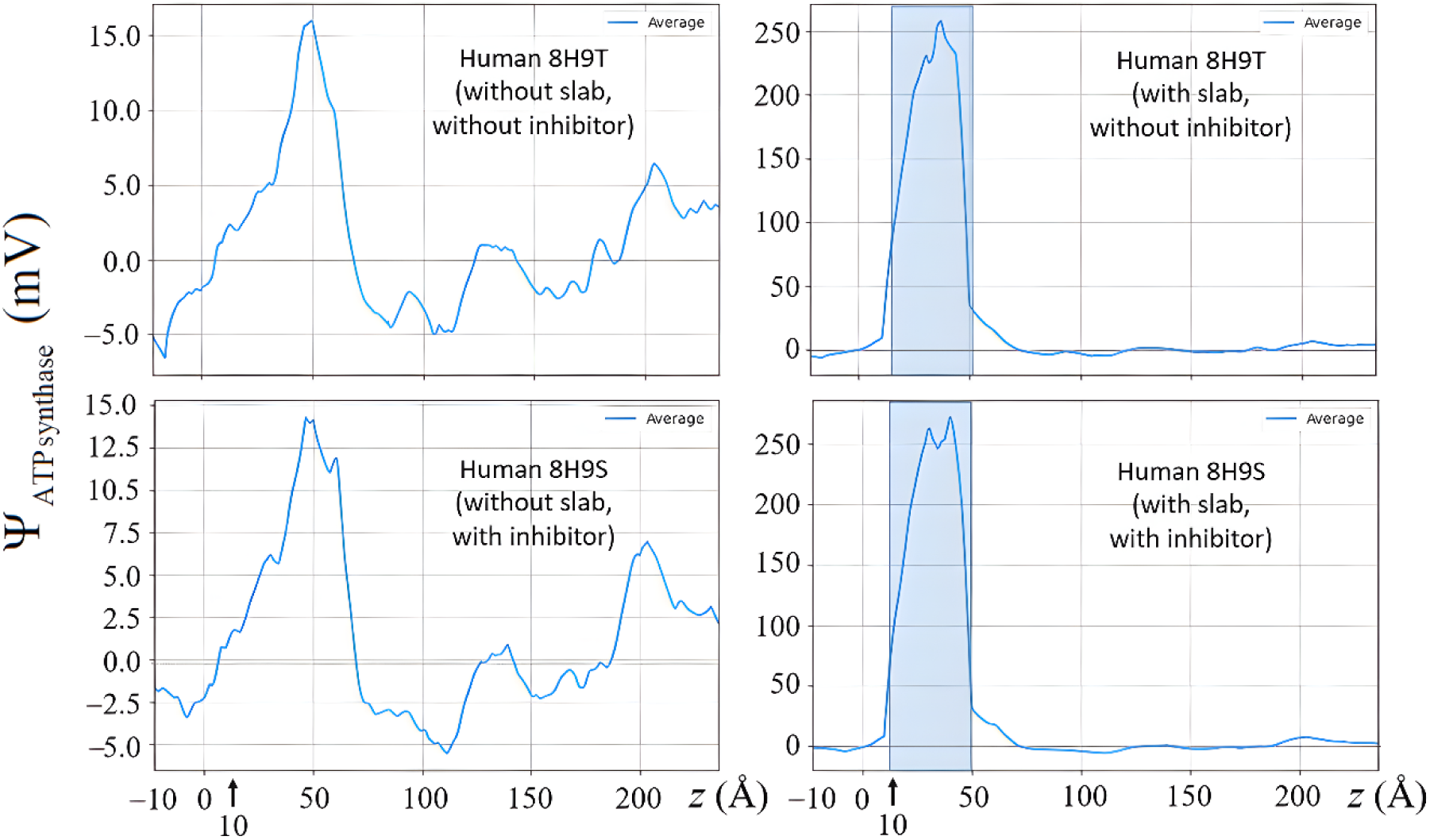
Axial electrostatic potential profiles for representative human ATP synthase structures with and without membrane embedding. **Left panels:** Plane-averaged electrostatic potential Ψ(*z*) (mV) along the long molecular axis (*z*) for PDB 8H9T (top, no inhibitor) and 8H9S (bottom, with endogenous inhibitor – see text) in fully aqueous medium (150 mM NaCl, no explicit membrane). Under these conditions, the magnitude of the potential difference between the exit-side and the entry-side windows fluctuates modestly (≈ ± 10–15 mV). **Right panels:** The same structures after embedding the F_o_ sector in a planar low-dielectric slab (ε = 4) spanning *z* = +10 to +50 Å (shaded region) to mimic the membrane core. Membrane insertion produces strong dielectric focusing within the slab, with Ψ(*z*) exceeding 250 mV before relaxing outside the membrane region. The enzyme-linked voltage used in Fig. 4 is obtained directly from these profiles as ΔΨ_ATP synthase_ = ⟨Ψ⟩_matrix_ (*z* = +50 Å to +70 Å) ― ⟨Ψ⟩_IMS_ (*z* = ―10Å to +10 Å)), see Eq. (3), defined analogously to the chemiosmotic membrane potential (Ψ_matrix_ ― Ψ_IMS_). Note the different *y*-axis scales: Aqueous profiles span approximately ±10 mV, whereas slab-embedded profiles span *ca*. >250 mV, highlighting the order-of-magnitude amplification introduced by the low membrane dielectric constant. The inhibitor induces only minor perturbations relative to this dominant dielectric effect.

Finally, note that the ordinate scales differ substantially between aqueous and slab-embedded profiles in Fig. 5: the aqueous plots span approximately ±15 mV, whereas the slab-embedded plots extend beyond 200 mV. This difference in scale emphasizes the order-of-magnitude enhancement of the enzyme-linked voltage arising from membrane dielectric contrast.

## 3. Energetic scale of the axial electrostatic descriptor

The potential differences reported here are static descriptors of the electrostatic landscape generated by the protein charge distribution under defined dielectric and ionic conditions. Nevertheless, the magnitude of the calculated axial potential differences raises a theoretical question: *if part of this protein-intrinsic electrostatic asymmetry is experienced by a proton during translocation, how large could its energetic effect be?* The following equations should therefore be read as a heuristic thermodynamic estimate, not as proof that ATP synthase generates an independent voltage source or that the pmf can be rigorously decomposed into two separately measurable voltages. Establishing this hypothesis will require pathway-resolved electrostatic sampling, molecular dynamics, constant-pH calculations, and mutational or biochemical validation. With these cautionary notes, one may define an effective local electrical potential difference experienced by the proton-translocation machinery as the sum of the chemiosmotic membrane contribution (ΔΨ_membrane_) and that of the protein itself (ΔΨ_ATP synthase_):

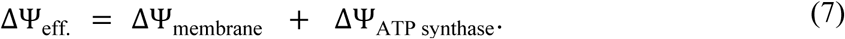

With this new heuristic term, Eq. (2) for the Gibbs energy released per proton may be written:

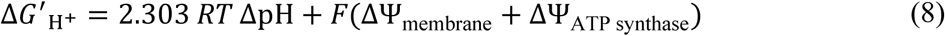

or equivalently,

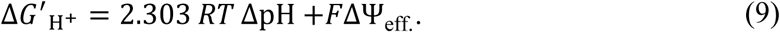

where Δ*G* _H+_ is the free energy associated with one proton translocation including the effect of the enzyme’s voltage – to be contrasted with Δ*G*_ATP_ which is the Gibbs energy of formation of one ATP molecule in cellular conditions independent of whether we account for the enzyme-linked voltage or not.

With this possible term, and from Eq. (8), the required membrane potential to drive ATP formation is:

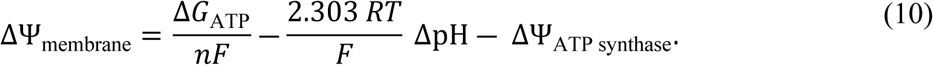

Hence, the sign of ΔΨ_ATP synthase_ determines whether the protein-intrinsic electrostatic asymmetry would, in principle, reduce or increase the membrane contribution needed to achieve a given ATP-synthesis free-energy demand. A contribution with the same effective axial electrostatic orientation as the chemiosmotic electrical term would be constructive, whereas one with the opposite polarity would be opposing. This asks whether protein electrostatics could locally bias how the electrical component of the electrochemical gradient is experienced by the F_o_ motor without any implication of creation or destruction of free energy by the protein.

To estimate the possible scale of the effect, consider Δ*G*_ATP_ ≈ 50 kJ·mol^−1^, *n* = 3 H^+^ per ATP, and ΔpH = 0.5. The chemical term contributes 2.303 *RT*ΔpH ≈ 3 kJ·mol^−1^ per proton (≈ 9 kJ·mol⁻¹ per ATP), leaving ∼41 kJ·mol^−1^ to be supplied electrically. Dividing by 3*F* yields a required combined electrical potential of ≈ 0.14 V. An enzyme-linked contribution of ± 20 mV would therefore shift the membrane potential requirement by ∼ 14% of this value. Importantly, *this does not imply the creation of additional free energy; rather, it redistributes the electrical component between the membrane dielectric and the intrinsic protein field*.

Depending on the metabolic pathway, the maximum P/O ratio (ATP produced per oxygen atom consumed) for an ATP synthase with 10 c-rings ranges from 1.385 to 2.494 [24]. When expressed per molecule of O₂ rather than per oxygen atom, these values should be doubled. A constructive enzyme-linked voltage would reduce the membrane contribution required per proton and could, in principle, slightly increase the attainable ATP yield under idealized, tightly coupled conditions. Conversely, a destructive contribution would increase the electrical demand on the membrane potential. In practice, experimentally observed P/O ratios are lower than theoretical maxima due to proton leak and other dissipative processes; thus, the enzyme-linked term should be interpreted as a modifier of electrical partitioning rather than a determinant of absolute bioenergetic efficiency.

## 4. Human ATP synthase displays an opposing axial electrostatic descriptor

Under the convention adopted here (ΔΨ_ATP synthase_ = ΔΨ_matrix_− ΔΨ_IMN_), the human enzyme exhibits a positive enzyme-linked voltage of approximately +0.02 V in the membrane-embedded model (Fig. 4b). Because the physiological chemiosmotic membrane potential during ATP synthesis is negative (ΔΨ_membrane_ < 0; matrix negative relative to IMS), the human ΔΨ_ATP synthase_ has the opposite sign and therefore partially offsets the membrane contribution. In this sense, the enzyme-linked field acts destructively with respect to the chemiosmotic driving force. The effective electrical term entering Eqs. (7) – (9) is therefore:

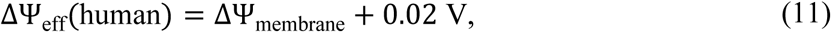

where ΔΨ_membrane_ < 0 under ATP-synthesis conditions. Thus, the magnitude of the total driving voltage is reduced by ∼20 mV relative to the membrane contribution alone.

The corresponding change in electrical free energy per proton is:

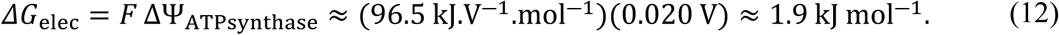

For *n* = 3 protons per ATP, this corresponds to ∼ 6 kJ·mol^−1^ less electrical work available for ATP formation; for *n* = 4, the shift is ∼ 8 kJ·mol^−1^.

Importantly, this does not imply the generation or loss of free energy by the enzyme itself; rather, it reflects redistribution of the electrical component of the proton-motive force between membrane dielectric and intrinsic protein electrostatics. In practical terms, a destructive enzyme-linked contribution increases the membrane potential magnitude required to reach a given Δ*G*_ATP_. In the illustrative example discussed above (requiring ∼ 0.14 V combined electrical potential), the membrane term must reach ∼ 0.16 V in magnitude if the enzyme contributes +0.02 V.

While the quantitative impact on steady-state ATP yield depends on coupling efficiency, proton leak, and other dissipative processes [13], a 10–20 mV intrinsic offset represents a thermodynamically non-negligible fraction of the electrical work per proton. Thus, even modest axial electrostatic asymmetries could, if sampled along the F_o_ proton pathway, influence local proton-transfer energetics. This remains a hypothesis requiring pathway-resolved and experimental validation.

## 5. Possible metabolic implications: a speculative translational outlook

This section is speculative and is included only to outline possible consequences that could be tested in future work. The static Poisson-Boltzmann descriptors reported here do not demonstrate altered proton flux, clinical nutrition requirements, body-mass regulation, or aging-related mitochondrial decline. These physiological quantities depend on rotary-state dynamics, protonation equilibria, lipid composition, proton leak, substrate supply, respiratory control, among several other factors. The purpose of this section is to identify where measurable consequences might arise if ATP-synthase electrostatic asymmetries are later shown to influence local coupling efficiency. Our purpose here is *not* to infer physiology from the present calculations.

The modified Weir equation (with protein correction) expresses the whole-organism *metabolic rate* (metabolic heat production (MPH)), in watts [35,36], commonly inferred from respiratory gas exchange using indirect calorimetry, as [37]:

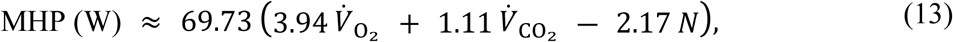

where *V*_O2_ and *V*_CO2_ are the rates of O_2_ consumption and of CO_2_ production (L·min⁻¹), and *N* is the urinary nitrogen excretion rate (g·min⁻¹), accounting for protein oxidation. In Eq. (13), the constants 3.94 and 1.11 have units kcal·L^⁻1^ for O_2_ and CO_2_, respectively; 2.17 has units kcal·g^⁻1^ for N; and 69.73 is the unit-conversion factor that converts the quantity in brackets from kcal·min^⁻1^ to watts. Although widely applied in human physiology, this formula is applicable to aerobic organisms with substrate corrections. This relationship reflects substrate oxidation chemistry and respiratory gas exchange. It is not modified by the present electrostatic calculations. A possible connection to ATP synthase electrostatics would enter only downstream, through the fraction of oxidation energy conserved as ATP versus dissipated as heat, and only if the electrostatic descriptors reported here are shown experimentally to affect coupling efficiency.

However, conversion of substrate oxidation into ATP depends on the effective electrical component of the proton-motive force. If enzyme-linked ESP modifies the voltage available to the F_o_ motor, the ATP synthesized per molecule of O_2_ reduced may shift slightly under tightly coupled conditions. To represent this effect, we may write a phenomenological correction to the classical P/O ratio:

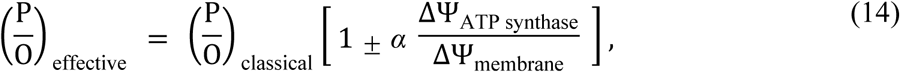

where 0 < *α* ≤ 1 is a dimensionless empirical coupling/attenuation factor that captures how strongly the enzyme-linked voltage perturbs the effective voltage drop experienced by the F_o_ motor. It represents the fraction of the intrinsic enzyme-linked field that effectively perturbs the voltage drop experienced by the F_o_ sector.

For representative mitochondrial values (|ΔΨ_membrane_| ≈ 150–180 mV), an intrinsic ± 20 mV offset constitutes a ∼ 10% perturbation of the electrical term. The caloric equivalent per liter O_2_ remains unchanged, but the fraction of oxidative energy conserved in ATP versus dissipated as heat may shift slightly if coupling is otherwise tight.

In practice, experimentally observed P/O ratios integrate additional processes such as proton leak, regulatory modulation, and structural variability [13]. The enzyme-linked voltage identified here should therefore be regarded as one contributor to energetic partitioning rather than a primary determinant of organismal metabolic rate. Animal physiology often notes that some species or conditions yield slightly different caloric equivalences. For instance, trained athletes can exhibit improved efficiency, burning slightly less O_2_ for the same mechanical work output. While some of that is due to biomechanical efficiency [38,39], it does not exclude that possibly a part of it could be traced to mitochondrial coupling (in)efficiency. These findings point to the possibility that species differences in ATP synthase’s electrostatic potential might underlie some variations in basal metabolic rate or thermogenesis.

Caloric equivalence and apparent metabolic economy can vary across species and physiological states. For example, trained individuals may consume slightly less O_2_ for a given mechanical output, partly due to biomechanical factors [38]. Although such differences are multifactorial, variation in mitochondrial coupling - including intrinsic electrostatic contributions of ATP synthase - could, in principle, contribute to differences in oxygen utilization or thermogenic balance.

Clinical nutrition commonly expresses substrate energy content using empirical caloric factors (e.g., 4 kcal·g^−1^ for carbohydrate, 9 kcal·g^−1^ for fat – the so-called Atwater factors used to estimate total parenteral nutrition (TPN)), reflecting the metabolizable enthalpy of oxidation after digestive and urinary corrections [40]. Implicit in this scheme is the assumption that mitochondrial coupling operates within a typical range, such that substrate oxidation yields a predictable ATP output:

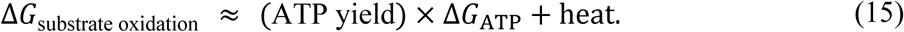

If the effective proton free energy is reduced by an opposing ΔΨ_ATP synthase_ < 0 then the ATP/O_2_ ratio is reduced. This means fewer ATP molecules are generated per mole of substrate oxidized even though the total heat release (kcal/g) remains unchanged. In practical terms, caloric adequacy and ATP-generating capacity need not be identical in individuals with reduced mitochondrial coupling efficiency. The present study does not show that ATP-synthase electrostatics measurably affects TPN response. Instead, it only identifies *one molecular variable* that could be examined in future studies of coupling efficiency.

While the discrepancy is modest in healthy individuals (on the order of ∼ 10%), in critically ill patients, neonates, cachectic patients, or those with mitochondrial disorders, the reduction in ATP yield per kcal could become clinically significant [41–43]. Whether this contributes to clinical observations such as failure to thrive despite apparent caloric sufficiency remains an open question requiring direct clinical and bioenergetic testing.

Could intrinsic electrostatic modulation of ATP synthase slightly alter the partitioning of dietary energy between ATP synthesis and heat, thereby influencing the relationship between caloric intake and net energy storage? This would occur *if and only if* the same caloric intake resulted in a slightly greater fraction of oxidation energy being dissipated as heat rather than conserved in ATP, holding behavior, appetite, and activity fixed. In this case, the net energy expenditure would be slightly higher hence, in humans for example, weight gain per labeled “calorie” could be slightly lower.

Quantitatively, the enzyme-linked offset is on the order of 10–20 mV, while the mitochondrial membrane potential magnitude is typically ∼ 150 mV, so the electrical driving term relevant to oxidative phosphorylation is perturbed at the level of roughly (10–20)/(150) ≈ 7–14%. However, only a fraction of whole-body energy balance is directly sensitive to this electrical component (because coupling is already influenced by proton leak, substrate cycling, variable P/O, and regulated thermogenesis), so the effective impact on total daily energy expenditure is plausibly much smaller.

We introduce an organism-level phenomenological attenuation factor κ that quantifies how much of a mitochondrial-scale perturbation in energy partitioning (here, an intrinsic ΔΨ_ATP synthase_ offset) is expressed in net whole-body energy balance after behavioral and physiological compensation. No direct empirical estimate of κ exists for this specific electrostatic mechanism; we therefore bracket its magnitude using two empirical constraints. First, whole-body energy-compensation analyses of imposed perturbations (e.g., exercise interventions) report compensation of ∼ 84–96%, corresponding to residual uncompensated fractions of ∼ 0.04–0.16 at the organism level [44]. This range provides a natural scale for the fraction of a mitochondrial energetic perturbation that could persist in net energy balance over time. Second, κ cannot plausibly exceed the fraction of oxidative energy routinely diverted to heat by mitochondrial inefficiencies such as proton leak; estimates of this diversion are ∼ 0.2–0.3 [45], which we treat as an upper physiological bound. Taking κ slightly above the lower residual-compensation range while remaining below this mitochondrial bound, we bracket κ ≈ 0.05–0.2.

As a purely illustrative upper-bound calculation, the product of a mitochondrial-scale electrical perturbation (0.06–0.13) and an organism-level attenuation factor κ (0.05–0.2) would yield approximately 0.3–2.6% of daily energy flux. For an intake of 2000 kcal·day^−1^, this corresponds to roughly 6–52 kcal·day^−1^. Using ∼ 7,700 kcal as the energy equivalent of 1 kg of adipose tissue [46], a persistent and uncompensated difference of 6 kcal·day^−1^ would amount to ∼ 1 kg over approximately 1,280 days (∼3.5 years), whereas 52 kcal·day^−1^ would correspond to ∼ 1 kg over ∼150 days (∼5 months). In practice, appetite regulation, spontaneous activity, endocrine responses, and adaptive thermogenesis would be expected to attenuate much of this theoretical imbalance, so any influence of intrinsic ATP synthase electrostatics would most plausibly manifest as a subtle shift in long-term energy partitioning rather than a major determinant of body mass trajectory.

Over long time scales, small differences in ATP/O_2_ yield could, in principle, interact with age-related changes in mitochondrial efficiency, but the present calculations do not establish such a mechanism. Mitochondrial coupling and oxidative phosphorylation capacity are known to decline with aging and in several chronic diseases, including metabolic syndrome and neurodegeneration, often accompanied by increased proton leak and altered respiratory control [47–50]. Because the intrinsic electrostatic profile of ATP synthase arises from the spatial distribution of charged residues and local dielectric environment, it is unlikely to be identical across tissues, physiological states, or individuals. Subtle, persistent differences in intrinsic ΔΨ_ATP synthase_, whether due to genetic variation, somatic mutation, oxidative damage, or altered membrane composition, could therefore contribute to small but cumulative shifts in ATP production efficiency and thermogenic balance over the human lifespan. We emphasize that this remains a testable hypothesis rather than a demonstrated mechanism; direct experimental quantification of ATP synthase electrostatic variation in aging and disease states will be required.

## 6. Limitations

Several limitations should be emphasized. First, the electrostatic potentials reported here are static continuum quantities obtained from Poisson–Boltzmann calculations on fixed protein structures; they do not describe explicit proton hopping, hydration dynamics, conformational fluctuations, or the time-dependent membrane potential. It is emphasized that the electrostatic potentials calculated in this work are not rotary-state-dependent. The c-ring has approximate *Cn* pseudosymmetry (e.g., *n* = 8 in *Homo sapiens*), whereas the central stalk and F_1_ sector have approximate *C*_3_ pseudosymmetry. This symmetry mismatch can result in phase-dependent electrostatic variations accompanying the rotation near the proton half-channels. The present axial profiles should therefore be read as static structural descriptors and not as time-averaged electrostatic fields over a full catalytic cycle.

Second, the plane-averaged potentials are global axial descriptors and should not be equated with the free-energy profile along the actual F_o_ proton-translocation pathway, which involves localized half-channels, protonatable residues, and transient hydrogen-bonded networks. Third, protonation states assigned by PROPKA at pH 7.0 are approximate and may differ under membrane potential, lipid-specific, or conformationally distinct conditions. Fourth, the membrane model used here is a simplified low-dielectric slab and does not include explicit lipid headgroups, cardiolipin enrichment, surface charge, membrane dipoles, or bilayer asymmetry. Thus, *the reported ΔΨ values should be interpreted as comparative, order-of-magnitude structure-derived electrostatic descriptors rather than independently measurable membrane voltages or demonstrated additive components of the proton-motive force*. Their functional relevance will require pathway-resolved electrostatic sampling, molecular dynamics, constant-pH calculations, mutagenesis, and direct biochemical or bioenergetic validation. Considerable further work is therefore needed before the proposed functional role of these electrostatic asymmetries can be firmly established.

## 7. Conclusions

We provide a comparative electrostatic analysis of 178 experimentally determined ATP synthase structures from 17 species. Across this dataset, ATP synthase exhibits reproducible species-dependent axial electrostatic asymmetries, with potential differences of the order of 10–20 mV between defined membrane-proximal axial windows in several taxa. These differences arise from the spatial distribution of charged residues, dielectric boundaries, and structural organization of the F_o_–F_1_ complex. They are static, structure-derived electrostatic descriptors rather than measurable membrane voltages or demonstrated additive components of the proton-motive force.

The observed profiles reveal substantial electrostatic diversity among ATP synthases. In particular, the five available *Homo sapiens* structures show a consistent axial electrostatic orientation within the human dataset. Broader variation is observed when comparing ATP synthases across the full set of 17 species. These observations reflect two distinct levels of comparison: within-species structural consistency for *Homo sapiens* and across-species electrostatic diversity for the broader dataset. The mycobacterial comparison is also noteworthy: *M. tuberculosis* shows a near-zero axial electrostatic asymmetry, whereas *M. smegmatis* trends more negative and is more variable. These differences may be relevant to future studies of proton-pathway energetics, species-specific coupling, and ATP-synthase inhibitors, but they do not by themselves establish altered proton flux, ATP yield, or drug sensitivity.

The possible functional significance of these electrostatic asymmetries remains a hypothesis to be confirmed by future more detailed investigations. If such protein-intrinsic potential differences persist near the F_o_ proton half-channels during the catalytic cycle, they could influence local protonation, proton release, proton transfer, or inhibitor binding. They should not, on the basis of the present static calculations alone, be treated as demonstrated additive components of the pmf. Testing this hypothesis will require pathway-resolved electrostatic calculations, molecular dynamics, constant-pH simulations, targeted mutagenesis of residues in the a-subunit and c-ring region, and biochemical measurements of ATP synthesis, proton flux, membrane potential, or inhibitor response.

More broadly, the present analysis shows that comparative protein electrostatics can reveal conserved and divergent features of large rotary enzymes that are not immediately apparent from structural coordinates alone. These results provide an electrostatic atlas of ATP synthase and a set of experimentally and computationally testable hypotheses concerning how protein charge architecture may shape local energy-transduction landscapes.

## Methods

### Dataset Collection

A total of 178 unique F_o_-F_1_-ATP synthase structures were retrieved from the Protein Data Bank (PDB), representing 17 distinct species. To our knowledge, this set encompasses the publicly available high-resolution structures of F-type ATP synthase at the time of data collection. Only experimentally solved structures were considered; predicted models (e.g., AlphaFold entries) were excluded. Structures were retained when the experimentally determined model was sufficiently complete to define the F_o_F_1_ complex, the c-ring/rotor axis, and the axial averaging windows reproducibly (entrance/IMS: *z* = ―10 to +10 Å, exit/matrix: *z* = +50 to +70 Å). No single hard resolution cutoff was imposed. Because continuum electrostatic calculations can be sensitive to local structural details, the reported values should be interpreted as comparative descriptors obtained under a uniform preparation protocol rather than as absolute potentials for individual structures.

### Workflow Overview

All computational analyses were performed using a Nextflow-based workflow (*ATPase_EP_Analyzer.nf*) developed in-house to ensure reproducibility and scalability on high-performance computing resources [51]. The workflow orchestrates multiple Python modules and external tools, managing structure preprocessing, coordinate alignment, electrostatic potential calculations, and data aggregation.

Because electrostatic descriptors of charged proteins (e.g., dipoles) are origin-dependent if they are extracted from the full Taylor expansion of the potential or field (as is commonly done), and as a good practice, our ESP analysis and cross-species comparisons of ΔΨATP_synthase_ were all performed in a consistent coordinate reference frame [52–53].

### Reproducibility

To facilitate reproducibility and to make the electrostatic protocol fully transparent, the main computational parameters used throughout the study are summarized in Table 1. These include the structural dataset, preparation workflow, protonation protocol, force-field assignment, dielectric constants, ionic conditions, membrane-slab model, coordinate-alignment convention, axial averaging windows, and software environment. The purpose of the table is to make explicit that all 178 ATP synthase structures were treated under the same electrostatic, geometric, and numerical assumptions, so that the reported differences reflect comparative structure-dependent trends rather than changes in computational setup.

**Table 1.**
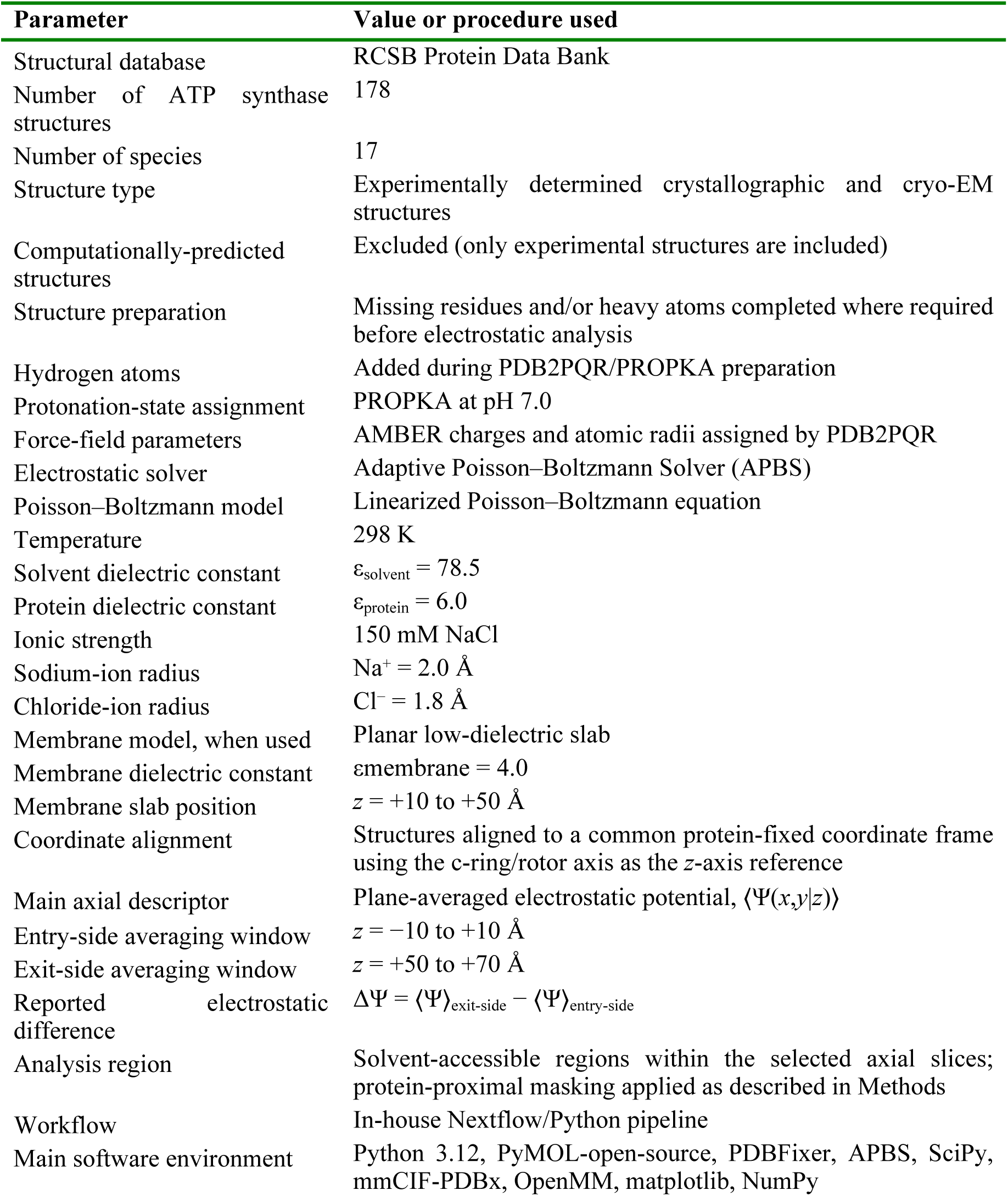
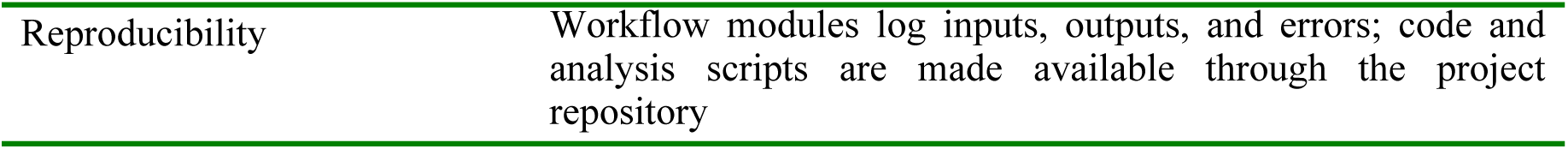
Computational parameters used for electrostatic-potential calculations.

The complete pipeline was executed under a Conda environment (Python 3.12, PyMOL-open-source, PDBFixer, APBS, SciPy, mmCIF-PDBx, OpenMM, matplotlib, NumPy). All steps were automated under Nextflow, enabling parallelization across multiple structures. Each module logs outputs and errors to ensure full reproducibility. (See Table 1).

### Structural Preprocessing and Alignment

1. Rotor (the c-ring) isolation and alignment were performed using a PyMOL-based script (*pymol_c_ring_aligner.py*), which automatically:

- Identifies the c-ring helices in each structure via MMCIF parsing,
- constructs pseudoatoms to define Cartesian reference axes,
- generates centroids (*centroid.py*) to approximate the c-ring central axis,
- aligns the rotor relative to the stator using pseudoatom vectors,
- outputs^1^ aligned whole ATP synthase and isolated c-ring PDB files for subsequent steps.
2. Protonation state assignment and electrostatic input generation were carried out with a wrapper script around PDB2PQR (*pdb2pqr.py*):

- Force field: AMBER.
- Protonation: PROPKA at physiological pH (7.0).
- Ion parameters: 150 mM NaCl (Na⁺ radius 2.0 Å, Cl⁻ radius 1.8 Å).
- Water dielectric constant: ε = 78.5.
- Protein dielectric constant: ε = 6.0.
3. APBS input files were automatically modified to include ion settings and standardized potential output (*\*-pot.dx*). For calculations with the membrane, a dielectric slab representing the inner mitochondrial membrane was added.

### Electrostatic Potential Calculation and Analysis

The resulting ESP maps were processed with the analysis module (*apbs_ctscan_analysis.py*), which implements the following procedure:

- Computation of the three-dimensional electrostatic potential Ψ_ATP synthase_(*x*,*y*,*z*) maps for each ATP synthase structure using APBS (*Adaptive Poisson–Boltzmann Solver*). APBS was executed on the PQR file with its custom “*.in*” parameters file generated by *pdb2pqr.py*.
- For calculations with the membrane, the generated maps are further modified to introduce a 40 Å membrane slab (ε = 4.0) dielectric model using *draw_membrane2a*, then the modified maps are resubmitted to APBS to perform the complete system simulation. The slab thickness is centered on the F_o_ rotor’s *z*-axis, spanning the rotor-stator interface from *z* = 10 Å to 50 Å.
- Protein geometric analysis is performed (*prot_dim*) to locate the geometric midpoint of the F_o_ rotor (*z*-axis reference).
- Slicing of the ESP grid (*ct_scan*) as a series of planar cross-sections perpendicular to the enzyme’s long (rotational) axis. Each section represents a two-dimensional map of Ψ_ATP synthase_(*x*,*y*) at a fixed *z*-coordinate, allowing direct inspection of the potential distribution along the proton-translocation pathway.
- A masking function (*mask_pot*) is implemented to restrict the analysis to solvent regions between 5 Å and 15 Å from the protein surface.
- The electric field is estimated by a numerical derivative (finite-difference) of the *z*-component of the electrostatic potential (*efield*).
- Angular averaging is applied (*ang_avg*) over circular slices to yield the mean ESP, standard deviations, and extrema as a function of *z*. Sectorized angular averages are also recorded.

The output of the procedure described above includes:

- Slice-averaged ESP curves (mean ± standard deviation).
- A set of planar sections of the Ψ_ATP synthase_(*x*,*y*) perpendicular to the long protein axis (z-axis) expressed numerically (two-dimensional grids) and visualized graphically for visual inspection.
- Summary data files for each structure containing *z*-averages (with their in-plane standard deviations). From the potential slices, potential differences (voltages) between intermembrane gap side and the matrix side regions are then estimated.

## Supporting Information

Python analysis notebook exported as PDF, showing the post-processing workflow for APBS-derived electrostatic-potential CSV files, including data loading, exclusion of non-F_o_F_1_ ATP synthase or otherwise unsuitable entries, classification of ATP synthase structures by axial electrostatic-potential profiles at the proton entry window, and generation of the combined and individual profile plots supporting the comparative electrostatic analysis.

## Acknowledgements

The authors thank Dahlia A. Awwad (Zewail City of Science and Technology) and Moustafa R. K. Ali (University of Nebraska Omaha) for helpful discussions. The authors are grateful to the financial support of the Natural Sciences and Engineering Council of Canada (NSERC), the Canadian Foundation for Innovation (CFI), Saint Mary’s University, Mount Saint Vincent University, the Digital Research Alliance of Canada for a DRI-EDIA Champions award to I.K.M., and for Research Nova Scotia for a Master’s Scotia Scholars Award (I.K.M).

## Declaration of generative AI and AI-assisted technologies in the manuscript preparation process

During the preparation of this work the authors used ChatGPT in order to condense and improve the writing structure in some passages. After using this tool/service, the authors reviewed and edited the content as needed and take full responsibility for the content of the published article.

## Footnotes

1 “Outputs” is used here as a verb.

