## Supplementary Information for "Comparative electrostatic mapping of ATP synthase reveals species-dependent axial asymmetry"

### SUPPORTING INFORMATION

**Description of the Supporting Information**

This Supporting Information file documents the post-processing and graphical analysis of the ATP synthase electrostatic-potential data used in the manuscript. It contains the following elements:

1. **Python analysis workflow**

   Code cells showing the Python libraries, data-loading functions, filtering routines, category definitions, and plotting functions used to process the APBS-derived electrostatic-potential CSV files.

2. **Data loading and preprocessing**

   Automated loading of PDB-derived electrostatic-potential CSV files, followed by exclusion of non-$F_oF_1$ ATP synthase or unsuitable entries before analysis.

3. **Classification of axial electrostatic profiles**

   Assignment of structures into three electrostatic-potential categories based on their axial plane-averaged profiles between $z$ = −20 to +20 Å: strong-positive, ambiguous or weak-positive, and negative categories.

4. **List of processed structures**

   Console output showing the PDB-derived CSV files successfully loaded into the analysis workflow.

5. **Summary of category counts**

   Printed output reporting the number of ATP synthase structures assigned to each electrostatic-potential category.

6. **Combined profile plots**

   Overlay plots of the one-dimensional axial electrostatic-potential profiles for each category, allowing comparison of the grouped profiles.

7. **Individual profile montages**

   Montage figures showing the individual axial electrostatic-potential profiles for structures in the negative, weak positive, and strong-positive categories.

8. **Computational provenance for figure generation**

   Documentation of the code and intermediate graphical outputs used to generate the comparative electrostatic-potential plots reported in the manuscript.

Together, these materials provide the computational provenance for the classification and visualization of the axial electrostatic-potential profiles used in the comparative analysis of ATP synthase.

In [41]:
```python
import os
import pandas as pd
import numpy as np
import matplotlib.pyplot as plt
from scipy.signal import find_peaks
from collections import defaultdict
```

Load electrostatic potential data from CSV files iteratively

In [42]:
```python
def load_csv_data(directory):
    """
    Loads data from all '_atpase_avg.csv' files in the given directory.
    Extracts data without headers and stores them in a dictionary with PDB codes as keys.
    """
    data_dict = {}

    for filename in os.listdir(directory):
        if filename.endswith("_atpase_avg.csv"):
            pdb_code = filename[:4]
            file_path = os.path.join(directory, filename)

            try:
                df = pd.read_csv(file_path)
                data_dict[pdb_code] = df
                print(f"Loaded: {filename}")
            except Exception as e:
                print(f"Error loading {filename}: {e}")

    return data_dict
```

Exclude data from multi-stator ATPases

In [43]:
```python
def exclude_prefixes(data_dict, prefixes=None):
    """
    Removes entries from data_dict whose keys start with any of the given prefixes.

    Parameters:
    data_dict (dict): Dictionary where keys are PDB codes and values are DataFrames.
    prefixes (set or list, optional): A set or list of prefixes to exclude. Defaults to a predefined set.

    Returns:
    dict: A new dictionary with the excluded entries removed.
    """
    if prefixes is None:
        prefixes = {
            "3j9t", "3j9u", "3j9v", "5gar", "5gas", "5tsj", "5vox", "5voy", "5voz",
            "5y5x", "5y5z", "5y60", "6o7v", "6o7w", "6o7x", "6qum", "6r0w", "6r0y",
            "6r0z", "6r10", "6vq6", "6vq7", "6vq8", "6xbw", "6xby", "7fda", "7fdb",
            "7fdc", "7khr", "7tmr", "7tms", "7tmt", "7unf", "7uw9", "7uwa", "7uwb",
            "7uwc", "7uwd", "9bra", "9brq", "9brr", "9brs", "9brt", "9bru"
        }

    filtered_dict = {key: value for key, value in data_dict.items() if key not in prefixes}

    print(f"Excluded {len(data_dict) - len(filtered_dict)} entries.")
    return filtered_dict
```

Analyze electrostatic potential peaks and valleys

In [44]:
```python
def format_pdb_codes(pdb_codes):
    """
    Formats PDB codes by grouping entries that share the first three letters.
    """
    grouped_codes = defaultdict(list)
    for pdb in pdb_codes:
        grouped_codes[pdb[:3]].append(pdb[3])

    formatted_output = []
    for prefix, suffixes in grouped_codes.items():
        formatted_output.append(f"{prefix} {', '.join(suffixes)}")

    return " | ".join(formatted_output)
```

In [45]:
```python
def analyze_peaks_and_valleys(data_dict, min_peak_prominence=0):
    """
    Analyzes electrostatic potential data and categorizes PDBs based on the first peak in the zero range (-20 to 20 Å).
    If no peak is found in this range, categorizes based on the average electrostatic potential within the zero range.
    Saves categorized results into separate CSV files.
    """
    category_negative = {}
    category_weak_positive = {}
    category_strong_positive = {}

    negative_data = []
```

```python
    weak_positive_data = []
    strong_positive_data = []

    for pdb_code, df in data_dict.items():
        if "Avg_Potential (mV)" not in df.columns:
            print(f"Skipping {pdb_code}: Missing expected column")
            continue

        z_coordinates = df["z_coordinate (Å)"].values
        potential_values = df["Avg_Potential (mV)"].values

        # Identify peaks and filter only positive peaks
        peaks, peak_props = find_peaks(potential_values, prominence=min_peak_prominence)
        peaks = [peak for peak in peaks if potential_values[peak] > 0]
        valleys, valley_props = find_peaks(-potential_values, prominence=min_peak_prominence)

        # Find first peak within zero range (-20 to 20 Å)
        zero_range_mask = (z_coordinates >= -20) & (z_coordinates <= 20)
        zero_range_peaks = [peak for peak in peaks if zero_range_mask[peak]]

        if zero_range_peaks:
            first_peak_idx = zero_range_peaks[0]
            first_peak_potential = potential_values[first_peak_idx]
        else:
            # No peak found in zero range, use the average electrostatic potential within the zero range
            zero_range_potentials = potential_values[zero_range_mask]
            if len(zero_range_potentials) == 0:
                print(f"Skipping {pdb_code}: No data in zero range")
                continue
            first_peak_potential = np.mean(zero_range_potentials)

        # Categorize based on the first peak in zero range or the average electrostatic potential in zero range
        if first_peak_potential < 0:
            category_negative[pdb_code] = df
            target_data = negative_data
        elif 0 <= first_peak_potential <= 10:
            category_weak_positive[pdb_code] = df
            target_data = weak_positive_data
        else:
            category_strong_positive[pdb_code] = df
            target_data = strong_positive_data

        # Store peaks and valleys in the appropriate category
        for i, peak in enumerate(peaks):
            target_data.append([pdb_code, "Peak", z_coordinates[peak], potential_values[peak], peak_props["prominences"][i]])
        for i, valley in enumerate(valleys):
            target_data.append([pdb_code, "Valley", z_coordinates[valley], potential_values[valley], -valley_props["prominence

    # Save categorized results to CSV
    pd.DataFrame(negative_data, columns=["PDB_Code", "Z_Coordinate", "Electrostatic_Potential", "Type", "Amplitude"]).to_csv("
    pd.DataFrame(weak_positive_data, columns=["PDB_Code", "Z_Coordinate", "Electrostatic_Potential", "Type", "Amplitude"]).to_
    pd.DataFrame(strong_positive_data, columns=["PDB_Code", "Z_Coordinate", "Electrostatic_Potential", "Type", "Amplitude"]).t

    # Print summary of categorized PDBs
    print(f"Number of PDBs categorized as 'Negative': {len(category_negative)}")
    print(format_pdb_codes(category_negative.keys()))

    print(f"Number of PDBs categorized as 'Weak Positive': {len(category_weak_positive)}")
    print(format_pdb_codes(category_weak_positive.keys()))

    print(f"Number of PDBs categorized as 'Strong Positive': {len(category_strong_positive)}")
    print(format_pdb_codes(category_strong_positive.keys()))

    return category_negative, category_weak_positive, category_strong_positive
```

###### Plot electrostatic potential signals

In [46]:

```python
def plot_electrostatic_potentials(category_negative, category_weak_positive, category_strong_positive):
    """
    Plots electrostatic potential values for three categories of PDBs: Negative, Weak Positive, and Strong Positive.
    Adds vertical lines at x = -10 (Proton Entry) and x = 70 (Proton Exit).
    """
    categories = {
        "Negative": category_negative,
        "Weak Positive": category_weak_positive,
        "Strong Positive": category_strong_positive
    }

    for category_name, category_data in categories.items():
        fig, ax = plt.subplots(figsize=(10, 6))

        # Plot each PDB's data
        for pdb_code, df in category_data.items():
            ax.plot(df["z_coordinate (Å)"], df["Avg_Potential (mV)"], label=pdb_code)

        # Add vertical lines
        #ax.axvline(x=-10, color='black', linestyle='--', label="Proton Entry (Average)")
```

```python
        #ax.axvline(x=70, color='black', linestyle='--', label="Proton Exit (Average)")

        # Labels and title
        ax.set_xlabel("Z Coordinate (Å)")
        ax.set_ylabel("Electrostatic Potential (mV)")
        ax.set_title(f"Electrostatic Potential Profiles ({category_name} Category)")

        # Legend settings
        ax.legend(loc='upper left', bbox_to_anchor=(1, 1), ncol=5)
        plt.grid()

        # Save and show
        plt.savefig(f"electrostatic_potential_{category_name.lower().replace(' ', '_')}.png", bbox_inches='tight')
        plt.show()
```

In [47]:
```python
def plot_montage(data_dict, category_name):
    """
    Plots each PDB in a separate panel to create a montage for better visualization.

    Parameters:
        data_dict (dict): Dictionary containing PDBs and their corresponding data.
        category_name (str): Name of the category (e.g., "Negative", "Weak Positive", "Strong Positive") to be used in the plo
    """
    num_pdbs = len(data_dict)
    if num_pdbs == 0:
        print(f"No PDBs available for {category_name} category.")
        return

    cols = min(5, num_pdbs)  # Define number of columns (max 5 per row)
    rows = (num_pdbs // cols) + (num_pdbs % cols > 0)  # Calculate necessary rows

    fig, axes = plt.subplots(rows, cols, figsize=(cols * 4, rows * 3), constrained_layout=True)
    axes = np.array(axes).flatten()  # Flatten in case of single row or column

    for ax, (pdb_code, df) in zip(axes, data_dict.items()):
        if "Avg_Potential (mV)" in df.columns and "z_coordinate (Å)" in df.columns:
            ax.plot(df["z_coordinate (Å)"], df["Avg_Potential (mV)"], label=pdb_code)
            ax.set_title(pdb_code)
            ax.set_xlabel("Z Coordinate (Å)")
            ax.set_ylabel("Electrostatic Potential (mV)")
            ax.grid()

    # Hide unused subplots if there are fewer PDBs than the grid size
    for i in range(num_pdbs, len(axes)):
        fig.delaxes(axes[i])

    plt.suptitle(f"Electrostatic Potential Montage - {category_name}", fontsize=14)
    plt.savefig(f"electrostatic_potential_montage_{category_name.replace(' ', '_')}.png", bbox_inches='tight')
    plt.show()
    print(f"Montage saved as: electrostatic_potential_montage_{category_name.replace(' ', '_')}.png")
```

In [48]:
```python
# Main
if __name__ == "__main__":
    csv_directory = "Pot_Avg_CSVs"
    data = load_csv_data(csv_directory)
    data = exclude_prefixes(data)
    min_prominence_threshold = 1.75
    zero_threshold = 20.0
    category_negative, category_weak_positive, category_strong_positive = analyze_peaks_and_valleys(data, min_peak_prominence=
    plot_electrostatic_potentials(category_negative, category_weak_positive, category_strong_positive)
    plot_montage(category_negative, "Negative")
    plot_montage(category_weak_positive, "Weak Positive")
    plot_montage(category_strong_positive, "Strong Positive")
```

```
Loaded: 2wpd_fixed_atpase_avg.csv
Loaded: 2xok_fixed_atpase_avg.csv
Loaded: 3j9t_fixed_atpase_avg.csv
Loaded: 3j9u_fixed_atpase_avg.csv
Loaded: 3j9v_fixed_atpase_avg.csv
Loaded: 5ara_fixed_atpase_avg.csv
Loaded: 5are_fixed_atpase_avg.csv
Loaded: 5arh_fixed_atpase_avg.csv
Loaded: 5ari_fixed_atpase_avg.csv
Loaded: 5dn6_fixed_atpase_avg.csv
Loaded: 5fij_fixed_atpase_avg.csv
Loaded: 5fik_fixed_atpase_avg.csv
Loaded: 5fil_fixed_atpase_avg.csv
Loaded: 5fl7_fixed_atpase_avg.csv
Loaded: 5gar_fixed_atpase_avg.csv
Loaded: 5gas_fixed_atpase_avg.csv
Loaded: 5lqx_fixed_atpase_avg.csv
Loaded: 5lqy_fixed_atpase_avg.csv
Loaded: 5lqz_fixed_atpase_avg.csv
Loaded: 5t4o_fixed_atpase_avg.csv
Loaded: 5t4p_fixed_atpase_avg.csv
Loaded: 5t4q_fixed_atpase_avg.csv
Loaded: 5tsj_fixed_atpase_avg.csv
Loaded: 5vox_fixed_atpase_avg.csv
Loaded: 5voy_fixed_atpase_avg.csv
Loaded: 5voz_fixed_atpase_avg.csv
Loaded: 5y5x_fixed_atpase_avg.csv
Loaded: 5y5z_fixed_atpase_avg.csv
Loaded: 5y60_fixed_atpase_avg.csv
Loaded: 6cp3_fixed_atpase_avg.csv
Loaded: 6cp6_fixed_atpase_avg.csv
Loaded: 6fkf_fixed_atpase_avg.csv
Loaded: 6fkh_fixed_atpase_avg.csv
Loaded: 6fki_fixed_atpase_avg.csv
Loaded: 6j5i_fixed_atpase_avg.csv
Loaded: 6j5j_fixed_atpase_avg.csv
Loaded: 6n2y_fixed_atpase_avg.csv
Loaded: 6n2z_fixed_atpase_avg.csv
Loaded: 6n30_fixed_atpase_avg.csv
Loaded: 6o7v_fixed_atpase_avg.csv
Loaded: 6o7w_fixed_atpase_avg.csv
Loaded: 6o7x_fixed_atpase_avg.csv
Loaded: 6oqr_fixed_atpase_avg.csv
Loaded: 6oqs_fixed_atpase_avg.csv
Loaded: 6oqt_fixed_atpase_avg.csv
Loaded: 6oqu_fixed_atpase_avg.csv
Loaded: 6oqv_fixed_atpase_avg.csv
Loaded: 6oqw_fixed_atpase_avg.csv
Loaded: 6pqv_fixed_atpase_avg.csv
Loaded: 6qum_fixed_atpase_avg.csv
Loaded: 6r0w_fixed_atpase_avg.csv
Loaded: 6r0y_fixed_atpase_avg.csv
Loaded: 6r0z_fixed_atpase_avg.csv
Loaded: 6r10_fixed_atpase_avg.csv
Loaded: 6rd9_fixed_atpase_avg.csv
Loaded: 6rdb_fixed_atpase_avg.csv
Loaded: 6rdc_fixed_atpase_avg.csv
Loaded: 6rde_fixed_atpase_avg.csv
Loaded: 6rdg_fixed_atpase_avg.csv
Loaded: 6rdh_fixed_atpase_avg.csv
Loaded: 6rdi_fixed_atpase_avg.csv
Loaded: 6rdj_fixed_atpase_avg.csv
Loaded: 6rdk_fixed_atpase_avg.csv
Loaded: 6rdl_fixed_atpase_avg.csv
Loaded: 6rdm_fixed_atpase_avg.csv
Loaded: 6rdn_fixed_atpase_avg.csv
Loaded: 6rdo_fixed_atpase_avg.csv
Loaded: 6rdp_fixed_atpase_avg.csv
Loaded: 6rdq_fixed_atpase_avg.csv
Loaded: 6rdr_fixed_atpase_avg.csv
Loaded: 6rds_fixed_atpase_avg.csv
Loaded: 6rdt_fixed_atpase_avg.csv
Loaded: 6rdu_fixed_atpase_avg.csv
Loaded: 6rdv_fixed_atpase_avg.csv
Loaded: 6rdw_fixed_atpase_avg.csv
Loaded: 6rdx_fixed_atpase_avg.csv
Loaded: 6rdy_fixed_atpase_avg.csv
Loaded: 6rdz_fixed_atpase_avg.csv
Loaded: 6re0_fixed_atpase_avg.csv
Loaded: 6re1_fixed_atpase_avg.csv
Loaded: 6re2_fixed_atpase_avg.csv
Loaded: 6re3_fixed_atpase_avg.csv
Loaded: 6re4_fixed_atpase_avg.csv
Loaded: 6re5_fixed_atpase_avg.csv
Loaded: 6re6_fixed_atpase_avg.csv
Loaded: 6re7_fixed_atpase_avg.csv
Loaded: 6re8_fixed_atpase_avg.csv
```

```
Loaded: 6re9_fixed_atpase_avg.csv
Loaded: 6rea_fixed_atpase_avg.csv
Loaded: 6reb_fixed_atpase_avg.csv
Loaded: 6rec_fixed_atpase_avg.csv
Loaded: 6red_fixed_atpase_avg.csv
Loaded: 6ree_fixed_atpase_avg.csv
Loaded: 6ref_fixed_atpase_avg.csv
Loaded: 6rep_fixed_atpase_avg.csv
Loaded: 6rer_fixed_atpase_avg.csv
Loaded: 6res_fixed_atpase_avg.csv
Loaded: 6ret_fixed_atpase_avg.csv
Loaded: 6reu_fixed_atpase_avg.csv
Loaded: 6tdy_fixed_atpase_avg.csv
Loaded: 6tdz_fixed_atpase_avg.csv
Loaded: 6te0_fixed_atpase_avg.csv
Loaded: 6tmh_fixed_atpase_avg.csv
Loaded: 6tt7_fixed_atpase_avg.csv
Loaded: 6vm1_fixed_atpase_avg.csv
Loaded: 6vm4_fixed_atpase_avg.csv
Loaded: 6vmb_fixed_atpase_avg.csv
Loaded: 6vmg_fixed_atpase_avg.csv
Loaded: 6vof_fixed_atpase_avg.csv
Loaded: 6voh_fixed_atpase_avg.csv
Loaded: 6voj_fixed_atpase_avg.csv
Loaded: 6vol_fixed_atpase_avg.csv
Loaded: 6von_fixed_atpase_avg.csv
Loaded: 6vq6_fixed_atpase_avg.csv
Loaded: 6vq7_fixed_atpase_avg.csv
Loaded: 6vq8_fixed_atpase_avg.csv
Loaded: 6wnq_fixed_atpase_avg.csv
Loaded: 6wnr_fixed_atpase_avg.csv
Loaded: 6xbw_fixed_atpase_avg.csv
Loaded: 6xby_fixed_atpase_avg.csv
Loaded: 6z1r_fixed_atpase_avg.csv
Loaded: 6z1u_fixed_atpase_avg.csv
Loaded: 6zpo_fixed_atpase_avg.csv
Loaded: 6zqm_fixed_atpase_avg.csv
Loaded: 6zqn_fixed_atpase_avg.csv
Loaded: 7fda_fixed_atpase_avg.csv
Loaded: 7fdb_fixed_atpase_avg.csv
Loaded: 7fdc_fixed_atpase_avg.csv
Loaded: 7jg5_fixed_atpase_avg.csv
Loaded: 7jg6_fixed_atpase_avg.csv
Loaded: 7jg7_fixed_atpase_avg.csv
Loaded: 7jg8_fixed_atpase_avg.csv
Loaded: 7jg9_fixed_atpase_avg.csv
Loaded: 7jga_fixed_atpase_avg.csv
Loaded: 7khr_fixed_atpase_avg.csv
Loaded: 7njk_fixed_atpase_avg.csv
Loaded: 7njl_fixed_atpase_avg.csv
Loaded: 7njm_fixed_atpase_avg.csv
Loaded: 7njn_fixed_atpase_avg.csv
Loaded: 7njo_fixed_atpase_avg.csv
Loaded: 7njp_fixed_atpase_avg.csv
Loaded: 7njq_fixed_atpase_avg.csv
Loaded: 7njr_fixed_atpase_avg.csv
Loaded: 7njs_fixed_atpase_avg.csv
Loaded: 7p2y_fixed_atpase_avg.csv
Loaded: 7p3n_fixed_atpase_avg.csv
Loaded: 7p3w_fixed_atpase_avg.csv
Loaded: 7tjy_fixed_atpase_avg.csv
Loaded: 7tjz_fixed_atpase_avg.csv
Loaded: 7tk0_fixed_atpase_avg.csv
Loaded: 7tk1_fixed_atpase_avg.csv
Loaded: 7tk2_fixed_atpase_avg.csv
Loaded: 7tk3_fixed_atpase_avg.csv
Loaded: 7tk4_fixed_atpase_avg.csv
Loaded: 7tk5_fixed_atpase_avg.csv
Loaded: 7tk6_fixed_atpase_avg.csv
Loaded: 7tk7_fixed_atpase_avg.csv
Loaded: 7tk8_fixed_atpase_avg.csv
Loaded: 7tk9_fixed_atpase_avg.csv
Loaded: 7tka_fixed_atpase_avg.csv
Loaded: 7tkb_fixed_atpase_avg.csv
Loaded: 7tkc_fixed_atpase_avg.csv
Loaded: 7tkd_fixed_atpase_avg.csv
Loaded: 7tke_fixed_atpase_avg.csv
Loaded: 7tkf_fixed_atpase_avg.csv
Loaded: 7tkg_fixed_atpase_avg.csv
Loaded: 7tkh_fixed_atpase_avg.csv
Loaded: 7tki_fixed_atpase_avg.csv
Loaded: 7tkj_fixed_atpase_avg.csv
Loaded: 7tkk_fixed_atpase_avg.csv
Loaded: 7tkl_fixed_atpase_avg.csv
Loaded: 7tkm_fixed_atpase_avg.csv
Loaded: 7tkn_fixed_atpase_avg.csv
Loaded: 7tko_fixed_atpase_avg.csv
```

```
Loaded: 7tkp_fixed_atpase_avg.csv
Loaded: 7tkq_fixed_atpase_avg.csv
Loaded: 7tkr_fixed_atpase_avg.csv
Loaded: 7tks_fixed_atpase_avg.csv
Loaded: 7tmr_fixed_atpase_avg.csv
Loaded: 7tms_fixed_atpase_avg.csv
Loaded: 7tmt_fixed_atpase_avg.csv
Loaded: 7unf_fixed_atpase_avg.csv
Loaded: 7uw9_fixed_atpase_avg.csv
Loaded: 7uwa_fixed_atpase_avg.csv
Loaded: 7uwb_fixed_atpase_avg.csv
Loaded: 7uwc_fixed_atpase_avg.csv
Loaded: 7uwd_fixed_atpase_avg.csv
Loaded: 7y5b_fixed_atpase_avg.csv
Loaded: 7y5c_fixed_atpase_avg.csv
Loaded: 7y5d_fixed_atpase_avg.csv
Loaded: 8dbp_fixed_atpase_avg.csv
Loaded: 8dbq_fixed_atpase_avg.csv
Loaded: 8dbr_fixed_atpase_avg.csv
Loaded: 8dbs_fixed_atpase_avg.csv
Loaded: 8dbt_fixed_atpase_avg.csv
Loaded: 8dbu_fixed_atpase_avg.csv
Loaded: 8dbv_fixed_atpase_avg.csv
Loaded: 8dbw_fixed_atpase_avg.csv
Loaded: 8f29_fixed_atpase_avg.csv
Loaded: 8f39_fixed_atpase_avg.csv
Loaded: 8fkj_fixed_atpase_avg.csv
Loaded: 8fl8_fixed_atpase_avg.csv
Loaded: 8g08_fixed_atpase_avg.csv
Loaded: 8g09_fixed_atpase_avg.csv
Loaded: 8g0a_fixed_atpase_avg.csv
Loaded: 8g0c_fixed_atpase_avg.csv
Loaded: 8g0d_fixed_atpase_avg.csv
Loaded: 8g0e_fixed_atpase_avg.csv
Loaded: 8h9s_fixed_atpase_avg.csv
Loaded: 8h9t_fixed_atpase_avg.csv
Loaded: 8h9u_fixed_atpase_avg.csv
Loaded: 8h9v_fixed_atpase_avg.csv
Loaded: 8j0s_fixed_atpase_avg.csv
Loaded: 8j0t_fixed_atpase_avg.csv
Loaded: 8jr0_fixed_atpase_avg.csv
Loaded: 8ki3_fixed_atpase_avg.csv
Loaded: 9bra_fixed_atpase_avg.csv
Loaded: 9brq_fixed_atpase_avg.csv
Loaded: 9brr_fixed_atpase_avg.csv
Loaded: 9brs_fixed_atpase_avg.csv
Loaded: 9brt_fixed_atpase_avg.csv
Loaded: 9bru_fixed_atpase_avg.csv
Excluded 44 entries.
Number of PDBs categorized as 'Negative': 35
6oq r, s, t, u, v, w | 6pq v | 6wn q, r | 6z1 r, u | 7nj k, l, m, n, o, p, q, s | 7p2 y | 7p3 n, w | 8db p, q, r, s, t, u, v, w
| 8h9 s, t, u, v | 8ki 3
Number of PDBs categorized as 'Weak Positive': 12
5t4 o, p, q | 6td y, z | 6te 0 | 6tt 7 | 6zp o | 6zq m, n | 7nj r | 8j0 s
Number of PDBs categorized as 'Strong Positive': 131
2wp d | 2xo k | 5ar a, e, h, i | 5dn 6 | 5fi j, k, l | 5fl 7 | 5lq x, y, z | 6cp 3, 6 | 6fk f, h, i | 6j5 i, j | 6n2 y, z | 6n3
0 | 6rd 9, b, c, e, g, h, i, j, k, l, m, n, o, p, q, r, s, t, u, v, w, x, y, z | 6re 0, 1, 2, 3, 4, 5, 6, 7, 8, 9, a, b, c, d,
e, f, p, r, s, t, u | 6tm h | 6vm 1, 4, b, g | 6vo f, h, j, l, n | 7jg 5, 6, 7, 8, 9, a | 7tj y, z | 7tk 0, 1, 2, 3, 4, 5, 6,
7, 8, 9, a, b, c, d, e, f, g, h, i, j, k, l, m, n, o, p, q, r, s | 7y5 b, c, d | 8f2 9 | 8f3 9 | 8fk j | 8fl 8 | 8g0 8, 9, a,
c, d, e | 8j0 t | 8jr 0
```

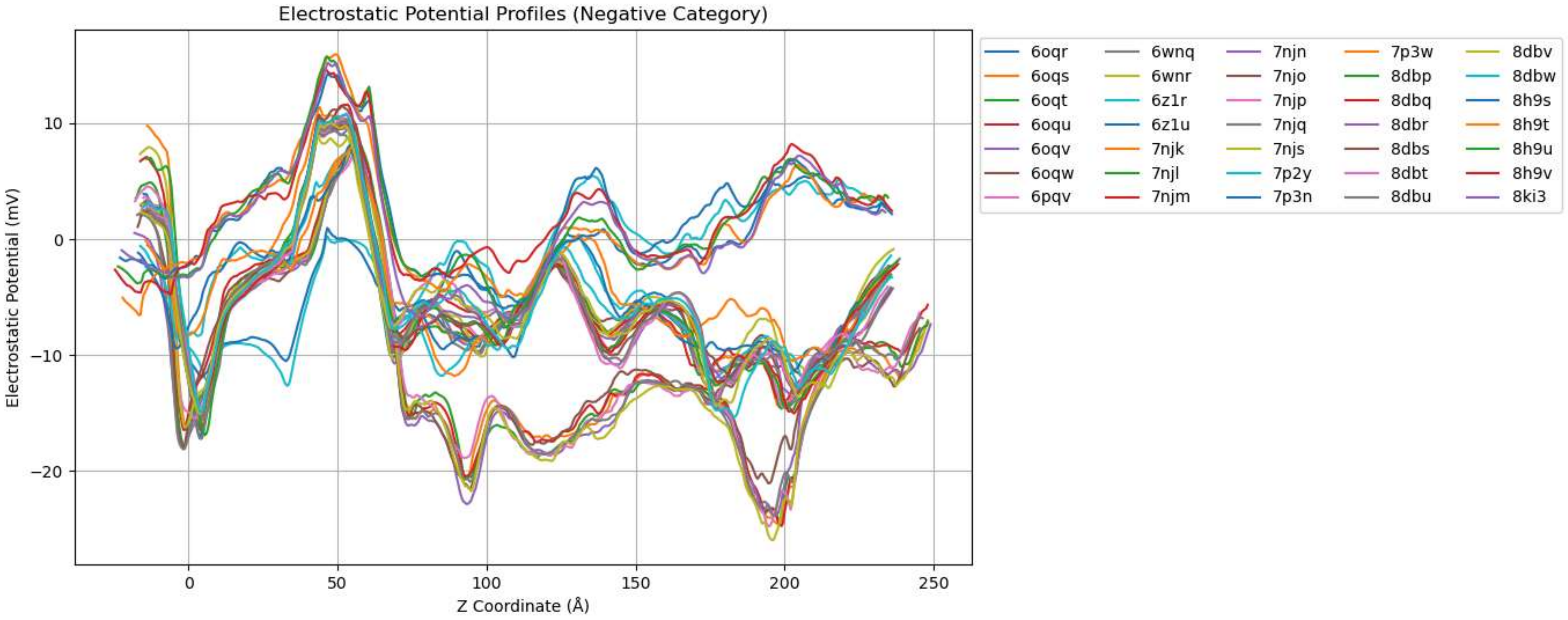

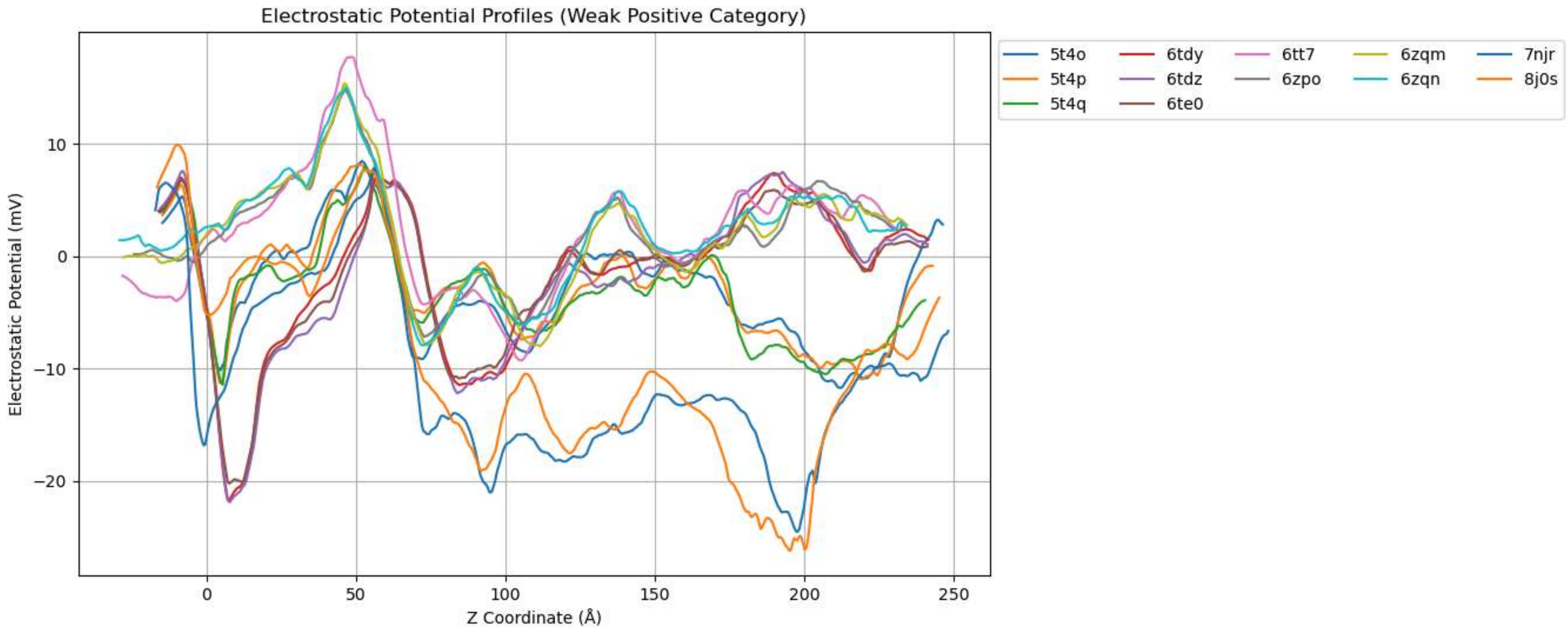
Electrostatic Potential Profiles (Weak Positive Category)
Electrostatic Potential (mV)
Z Coordinate (Å)
5t4o
5t4p
5t4q
6tdy
6tdz
6te0
6tt7
6zpo
6zqm
6zqn
7njr
8j0s

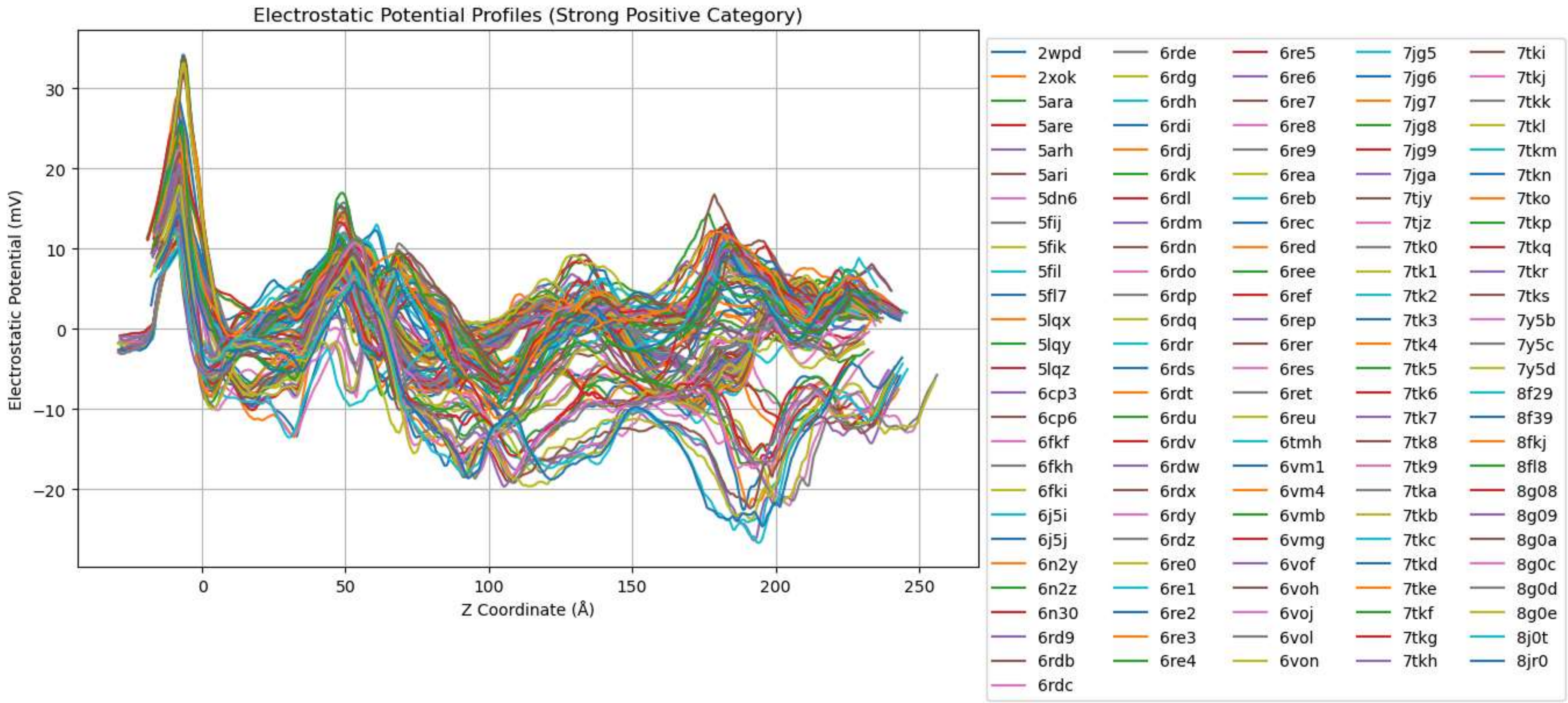
Electrostatic Potential Profiles (Strong Positive Category)
Electrostatic Potential (mV)
Z Coordinate (Å)
2wpd
2xok
5ara
5are
5arh
5ari
5dn6
5fij
5fik
5fil
5fl7
5lqx
5lqy
5lqz
6cp3
6cp6
6fkf
6fkh
6fki
6j5i
6j5j
6n2y
6n2z
6n30
6rd9
6rdb
6rdc
6rde
6rdg
6rdh
6rdi
6rdj
6rdk
6rdl
6rdm
6rdn
6rdo
6rdp
6rdq
6rdr
6rds
6rdt
6rdu
6rdv
6rdw
6rdx
6rdy
6rdz
6re0
6re1
6re2
6re3
6re4
6re5
6re6
6re7
6re8
6re9
6rea
6reb
6rec
6red
6ree
6ref
6rep
6rer
6res
6ret
6reu
6tmh
6vm1
6vm4
6vmb
6vmg
6vof
6voh
6voj
6vol
6von
7jg5
7jg6
7jg7
7jg8
7jg9
7jga
7tjy
7tjz
7tk0
7tk1
7tk2
7tk3
7tk4
7tk5
7tk6
7tk7
7tk8
7tk9
7tka
7tkb
7tkc
7tkd
7tke
7tkf
7tkg
7tkh
7tki
7tkj
7tkk
7tkl
7tkm
7tkn
7tko
7tkp
7tkq
7tkr
7tks
7y5b
7y5c
7y5d
8f29
8f39
8fkj
8fl8
8g08
8g09
8g0a
8g0c
8g0d
8g0e
8j0t
8jr0

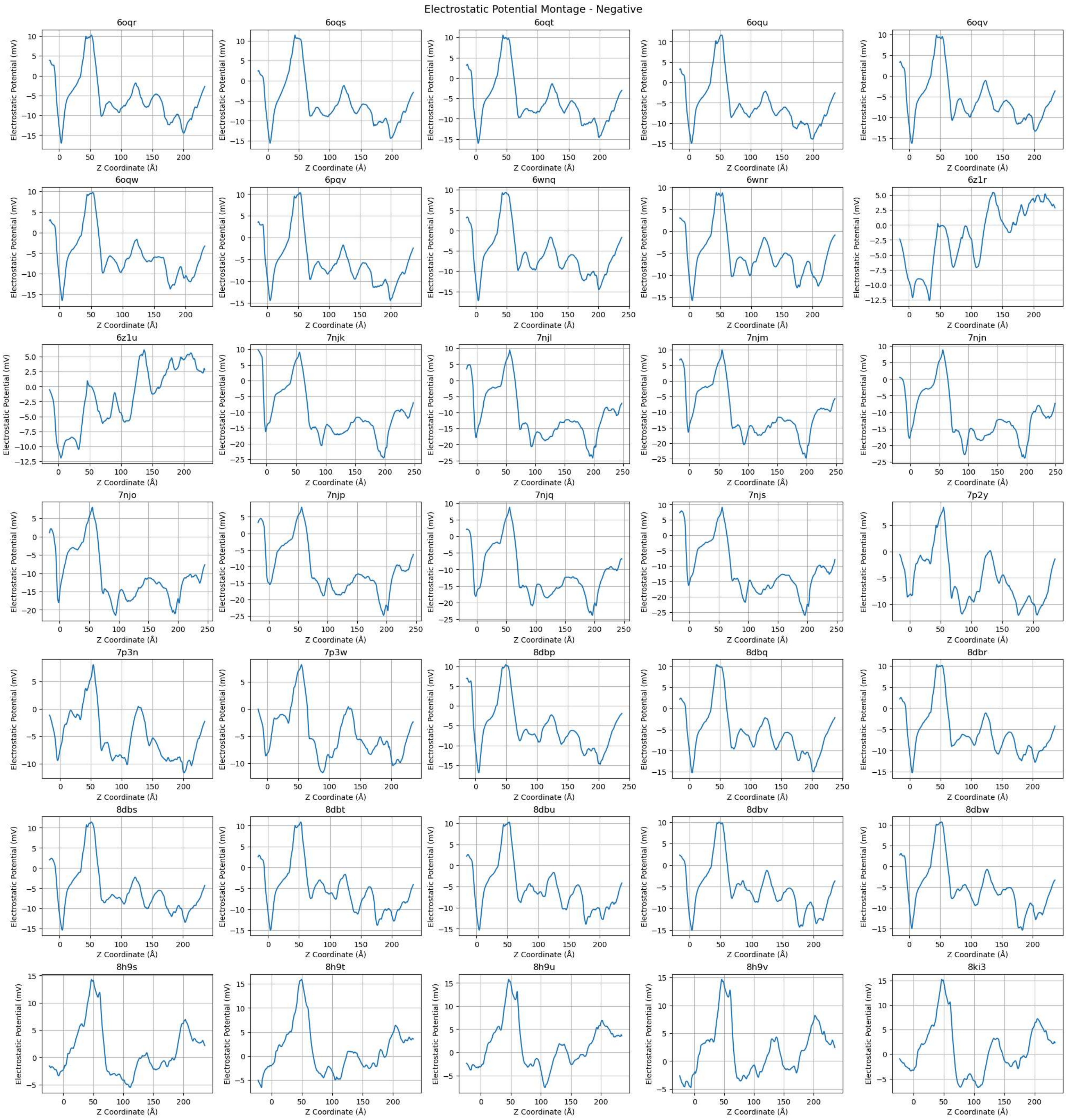


Montage saved as: electrostatic_potential_montage_Negative.png

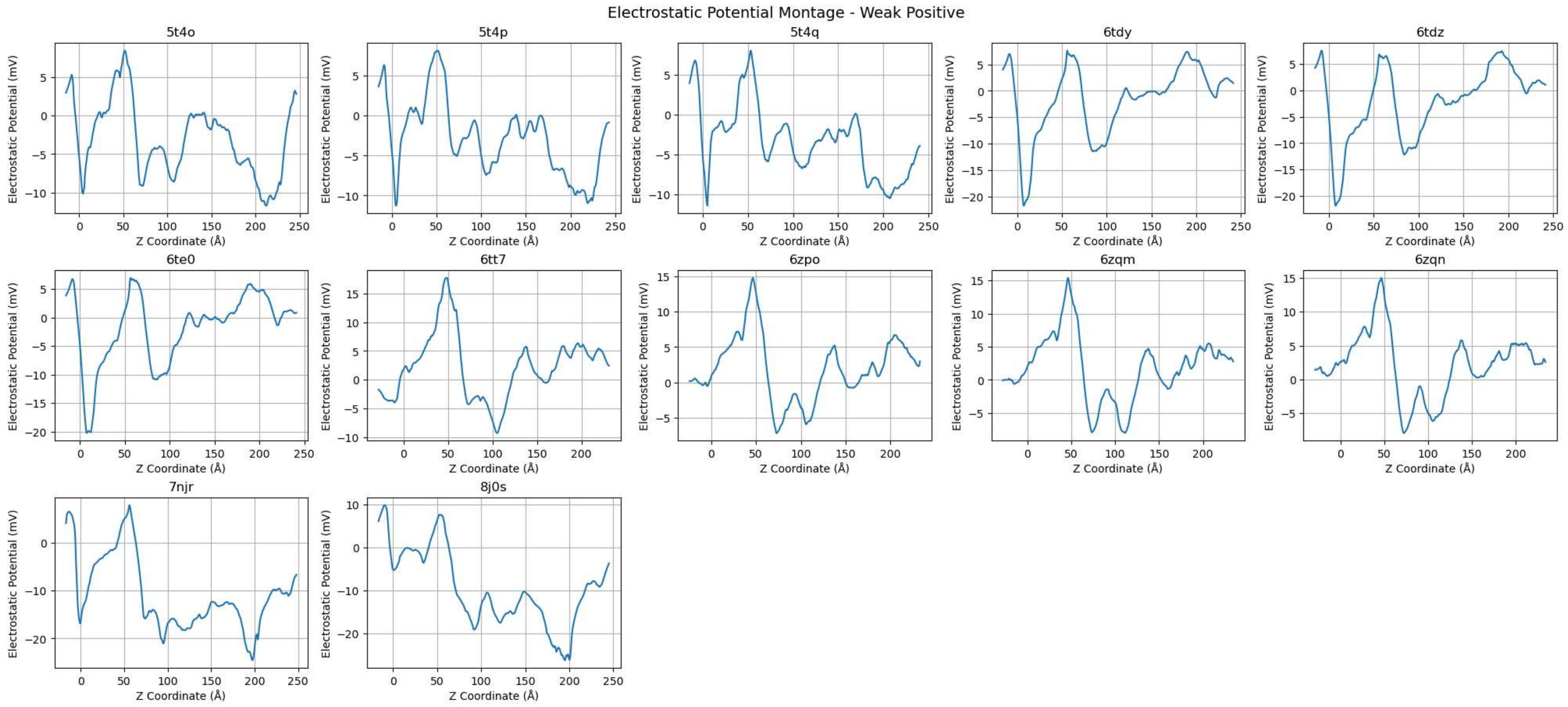


Montage saved as: electrostatic_potential_montage_Weak_Positive.png

Electrostatic Potential Montage - Strong Positive

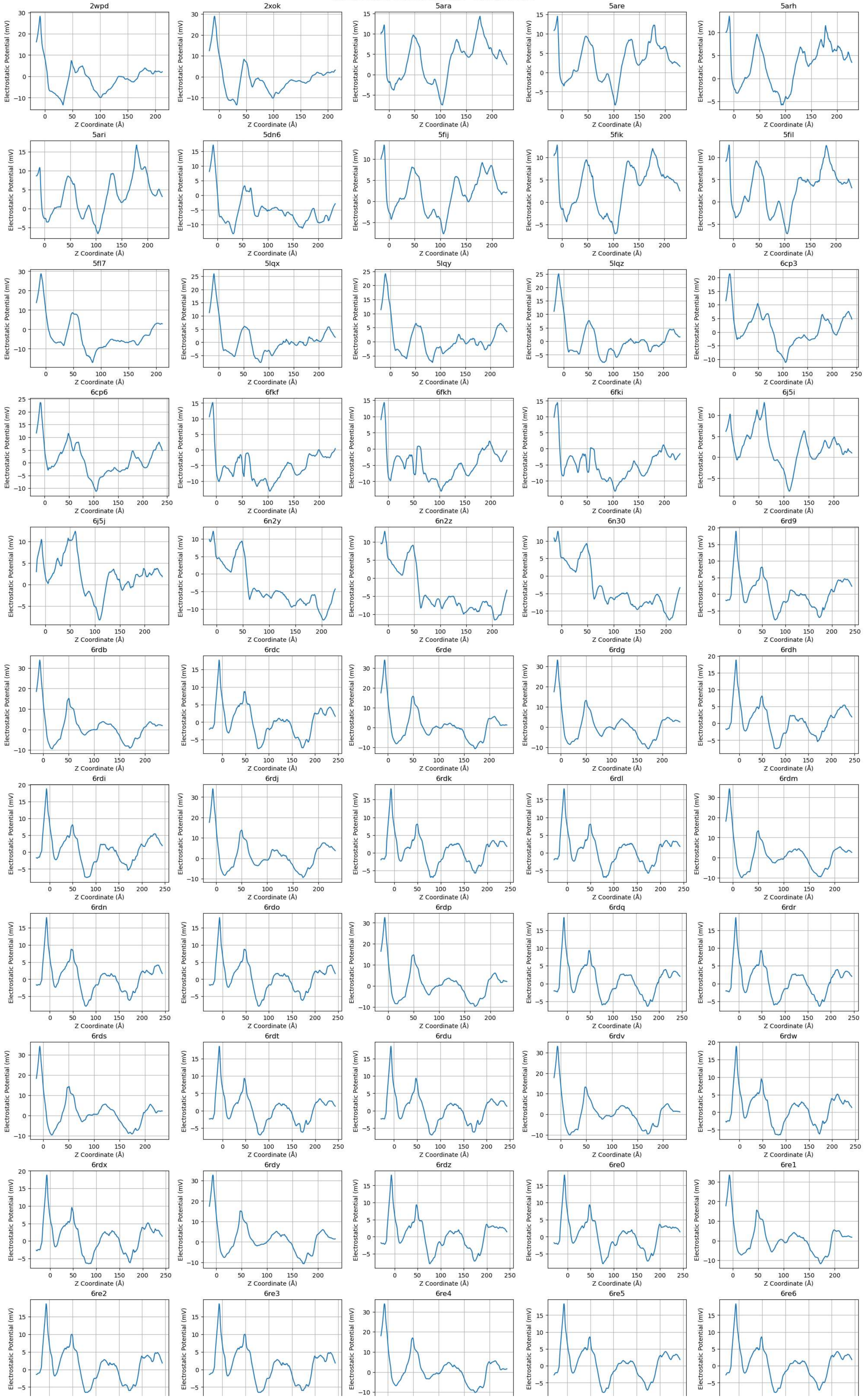

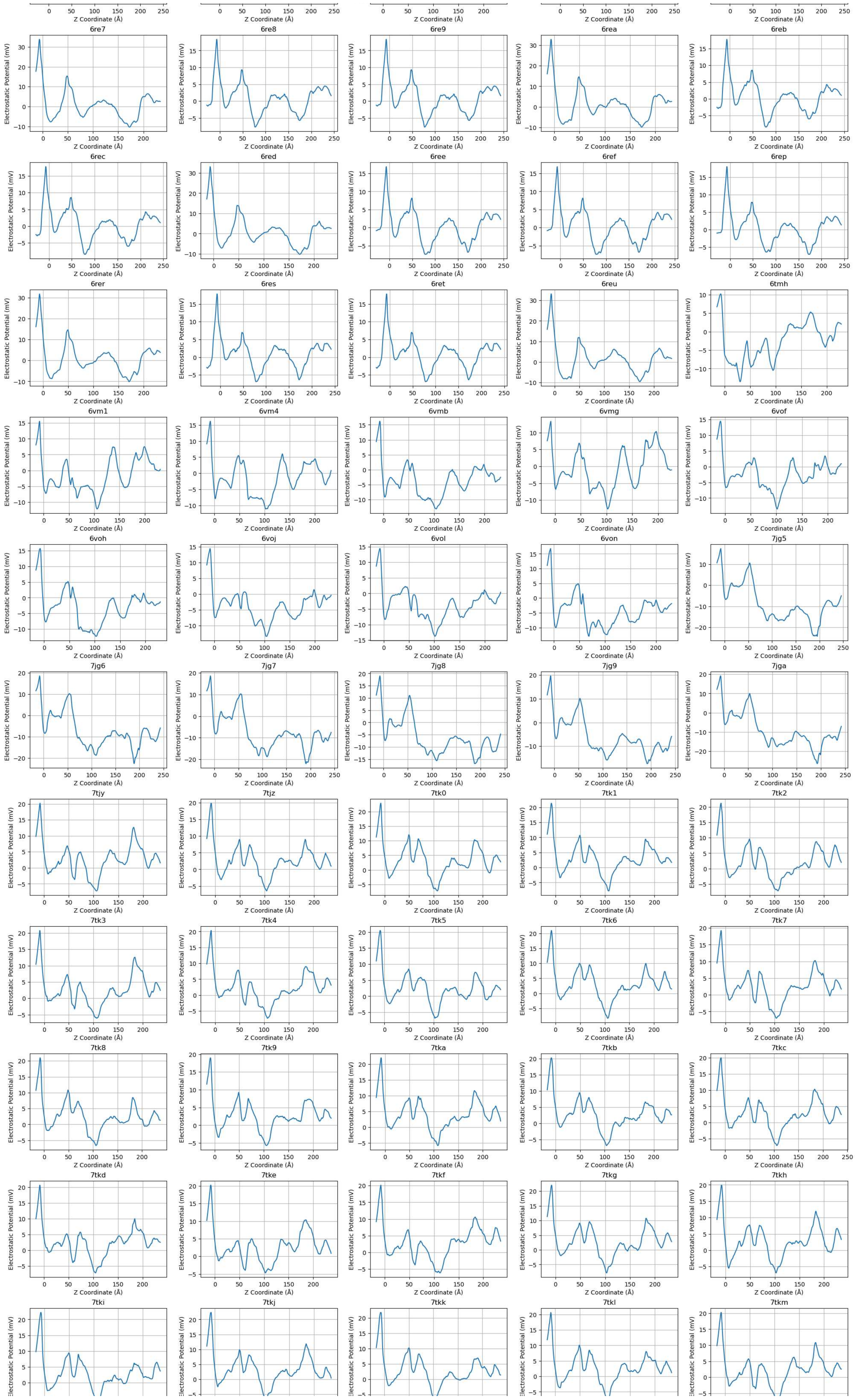

Z Coordinate (Å)
Electrostatic Potential (mV)
6re7
6re8
6re9
6rea
6reb
6rec
6red
6ree
6ref
6rep
6rer
6res
6ret
6reu
6tmh
6vm1
6vm4
6vmb
6vmg
6vof
6voh
6voj
6vol
6von
7jg5
7jg6
7jg7
7jg8
7jg9
7jga
7tjy
7tjz
7tk0
7tk1
7tk2
7tk3
7tk4
7tk5
7tk6
7tk7
7tk8
7tk9
7tka
7tkb
7tkc
7tkd
7tke
7tkf
7tkg
7tkh
7tki
7tkj
7tkk
7tkl
7tkm

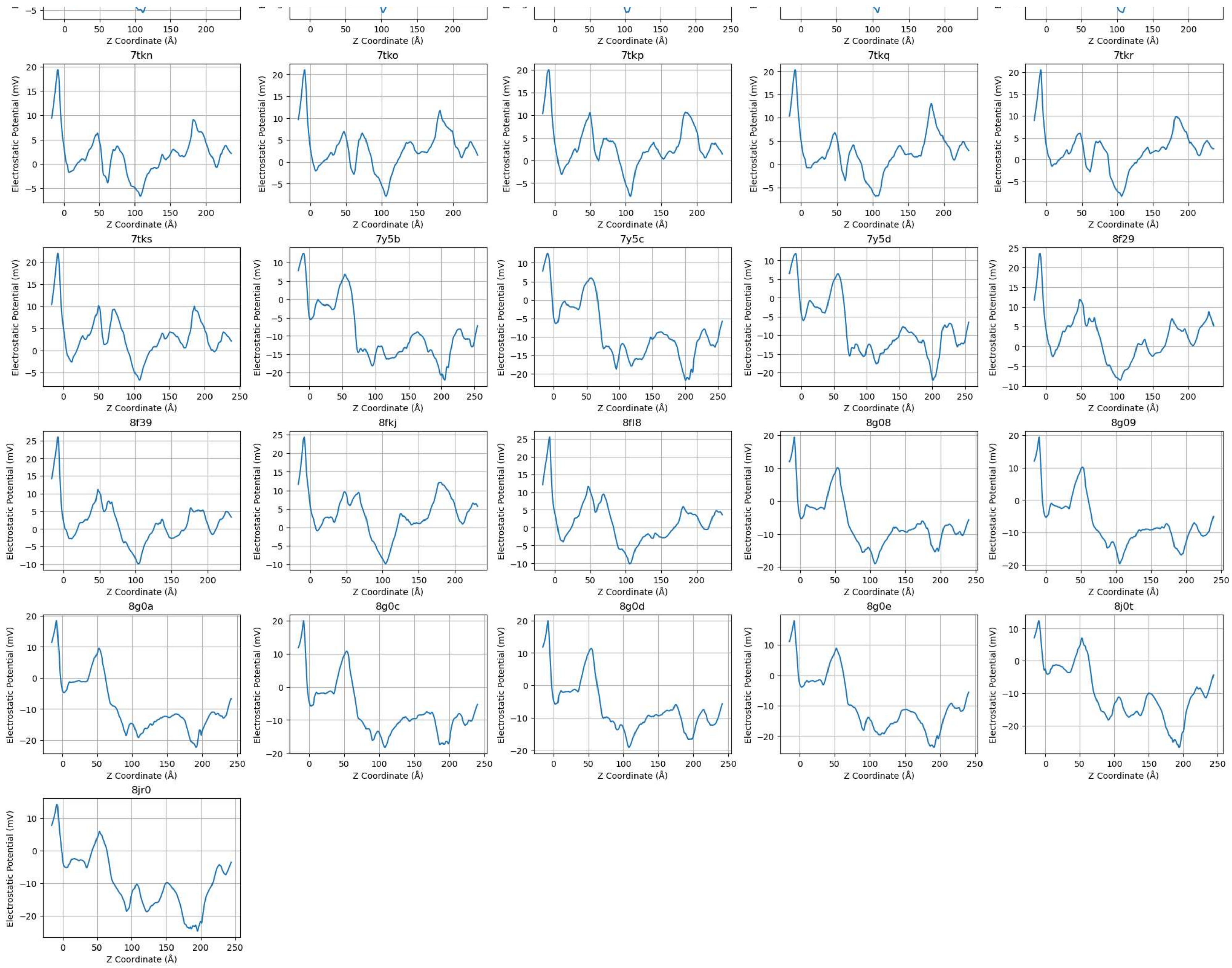


Montage saved as: electrostatic_potential_montage_Strong_Positive.png
